# Direct mitochondrial import of lactate supports resilient carbohydrate oxidation

**DOI:** 10.1101/2024.10.07.617073

**Authors:** Ahmad A. Cluntun, Joseph R. Visker, Jesse N. Velasco-Silva, Marisa J. Lang, Luis Cedeño-Rosario, Thirupura S. Shankar, Rana Hamouche, Jing Ling, Ji Eon Kim, Ashish G. Toshniwal, Hayden K. Low, Corey N. Cunningham, James Carrington, Jonathan Leon Catrow, Quentinn Pearce, Mi-Young Jeong, Alex J. Bott, Álvaro J. Narbona-Pérez, Claire E. Stanley, Qing Li, David R. Eberhardt, Jeffrey T. Morgan, Tarun Yadav, Chloe E. Wells, Dinesh K. A. Ramadurai, Wojciech I. Swiatek, Dipayan Chaudhuri, Jeffery D. Rothstein, Deborah M. Muoio, Joao A. Paulo, Steven P. Gygi, Steven A. Baker, Sutip Navankasattusas, James E. Cox, Katsuhiko Funai, Stavros G. Drakos, Jared Rutter, Gregory S. Ducker

**Author notes:** Correspondence and requests for materials should be addressed to G.S.D., or to S.G.D., or to J.R. These authors contributed equally to this work (Ahmad A. Cluntun, Joseph R. Visker, Jesse N. Velasco-Silva). Department of Biochemistry and Molecular Biology, Rutgers University, Piscataway, NJ 08854.

## Abstract

Lactate is the highest turnover circulating metabolite in mammals. While traditionally viewed as a waste product, lactate is an important energy source for many organs, but first must be oxidized to pyruvate for entry into the tricarboxylic acid cycle (TCA cycle). This reaction is thought to occur in the cytosol, with pyruvate subsequently transported into mitochondria via the mitochondrial pyruvate carrier (MPC). Using ^13^C stable isotope tracing, we demonstrated that lactate is oxidized in the myocardial tissue of mice even when the MPC is genetically deleted. This MPC-independent lactate import and mitochondrial oxidation is dependent upon the monocarboxylate transporter 1 (MCT1/*Slc16a1*). Mitochondria isolated from the myocardium without MCT1 exhibit a specific defect in mitochondrial lactate, but not pyruvate, metabolism. The import and subsequent mitochondrial oxidation of lactate by mitochondrial lactate dehydrogenase (LDH) acts as an electron shuttle, generating sufficient NADH to support respiration even when the TCA cycle is disrupted. In response to diverse cardiac insults, animals with hearts lacking MCT1 undergo rapid progression to heart failure with reduced ejection fraction. Thus, the mitochondrial import and oxidation of lactate enables carbohydrate entry into the TCA cycle to sustain cardiac energetics and maintain myocardial structure and function under stress conditions.

## Main

The oxidation of carbon fuels (e.g. carbohydrates and lipids) within the mitochondria is responsible for the generation of over 95% of ATP in humans^1,2^. Dietary sugars are the largest macronutrient in the Western diet, accounting for approximately 45% percent of energy intake^3^. Glucose generates ATP anaerobically by glycolysis to make pyruvate and aerobically upon the burning of pyruvate in the tricarboxylic acid (TCA) cycle^4^. To sustain glycolytic flux and ATP production, cytosolic NAD^+^ must be regenerated; this is achieved either by cytosol to mitochondria electron shuttles when pyruvate is oxidized within mitochondria or by the reduction of pyruvate to lactate in the cytosol by lactate dehydrogenase (LDH). Measurements showing that the flux of lactate is twice that of glucose within mammalian circulation suggests that lactate production is the dominant mode of NAD^+^ regeneration^5,6^.

High circulating lactate also serves as a major oxidative fuel source in both fasted and fed mammalian physiology^7–9^. Biochemically, the oxidation of lactate is thought to commence via the generation of cytosolic pyruvate, which is imported into the mitochondria by the obligate heterodimeric mitochondrial pyruvate carrier (MPC, encoded by *Mpc1* and *Mpc2*)^10,11^. This cytosolic LDH-dependent scenario leads to competition between LDH and glycolytic enzymes for NAD+, restricting flux through these essential pathways when metabolic demand is high (Fig. 1a). However, the heart is well-described as being capable of dynamically increasing the oxidation of lactate and glucose simultaneously in response to energetic demand^12–14^, implying a separation between glycolytic pyruvate production and lactate oxidation^15,16^. Here we reconcile the conflict between lactate, glucose and pyruvate oxidation in the myocardium by demonstrating that the lactate monocarboxylate transporter 1 (MCT1), which is canonically on the plasma membrane, is also present on the mitochondrial inner membrane. MCT1 is necessary for the import and oxidation of lactate and provides essential redundancy for heart energy metabolism such that when absent, hearts are unable to compensate in response to cardiac insults, leading to significant myocardial structural and functional impairment and heart failure.

**Fig. 1.**
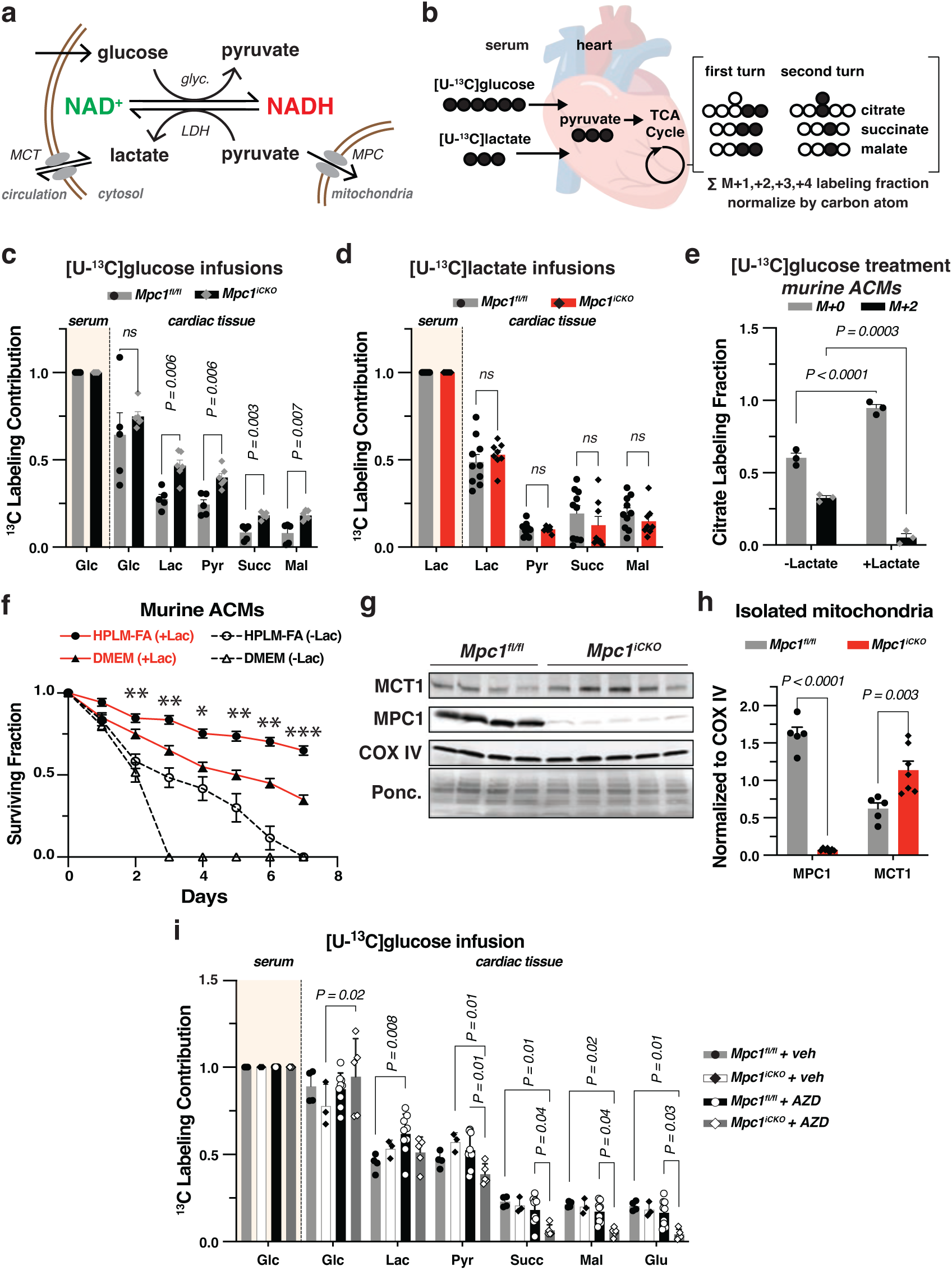
Lactate contributes to the TCA cycle independent of the MPC. **a,** Schematic representation of the competition for cytosolic NAD+ between glycolysis and lactate consumption, illustrating that rRedox balanced metabolism is lactate producing. Glyc. denotes glycolysis. **b**, Schematic of carbon labeling of heart TCA cycle metabolites resulting from infusion of [U-^13^C]glucose or [U-^13^C]lactate. **c**, Heart ^13^C labeling of lactate, pyruvate and TCA cycle metabolites following [U-^13^C]glucose infusion. **d**, Heart ^13^C metabolite labeling following [U-^13^C]lactate infusion in *Mpc1^iCKO^* (*n=10*) and *Mpc1^fl/fl^* (*n=8*). **e**, ^13^C enrichment of citrate from primary cultured adult cardiomyocytes (ACMs) cultured with [U-^13^C]glucose with or without 1.6 mM lactate, *n=3*. **f**, Survival fraction of ACMs grown in DMEM and HPLM-FA medias with and without lactate, *n=3-4* ACM preparations from distinct hearts. ^∗^p < 0.05, ^∗∗^p < 0.01, ^∗∗∗^p < 0.001 for HPLM-FA with and without lactate by unpaired t-test corrected for multiple comparisons at FDR 0.01. **g**, Immunoblots of MCT1 and MPC1 from purified mitochondria isolated from *Mpc1^fl/fl^* and *Mpc1^iCKO^* murine hearts. **h**, Quantification of MPC1 and MCT1 protein levels (*Mpc1^fl/fl^*, *n=5*, *Mpc1^iCKO^*, *n*=7). **l**, Heart ^13^C labeling of glycolytic and TCA cycle metabolites from *Mpc1^fl/fl^* and *Mpc1^iCKO^* mice treated with either AZD3965 (AZD) or vehicle (Veh) and infused with [U-^13^C]glucose (*Mpc1^fl/fl^*Veh *n=4*, *Mpc1^iCKO^* Veh *n=3*, *Mpc1^fl/fl^* AZD *n=8*, *Mpc1^iCKO^ n*=5). Glc: glucose; Lac: lactate; Pyr: pyruvate; Succ: succinate; Mal: Malate. All data represent mean ± SEM. Significance determined by multiple comparisons corrected (Holm-Sídák method) unpaired t tests (c and d), one-way ANOVA with Dunnett’s multiple comparison test (f, h, and k), and two-way ANOVA corrected for multiple comparisons (l). ns, not significant (p>0.05).

### Glucose carbon can enter the mitochondria independent of the MPC

To test the prevailing model of lactate oxidation, we employed our previously described tamoxifen-inducible cardiac-specific *Mpc1* knockout mice (*Mpc1^fl/fl^*:αMHC^MerCreMer*+/-*^, hereafter *Mpc1^iCKO^*) to quantify the extent to which the oxidation of glucose and lactate in the myocardium requires the MPC^17^. We implanted jugular vein catheters into 12-week old *Mpc1^iCKO^* and littermate controls (*Mpc1^fl/fl^*) with normal cardiac function, and subsequently infused awake animals with [U-^13^C]glucose or [U-^13^C]lactate to reach stable serum enrichment (Extended Data Fig. 1a). Hearts and serum were then harvested for isotopic labeling analysis of metabolites by liquid chromatography-mass spectrometry (LC-MS) (Fig. 1b). Similar to prior reports, we observed rapid appearance of ^13^C-lactate in serum upon [U-^13^C]glucose infusion, and vice versa with [U-^13^C]lactate infusion, reflecting high rates of interconversion between these metabolites (Extended Data Fig. 1b,c)^6^. Total isotopic enrichment and endogenous rates of glucose and lactate appearance were not different between *Mpc1^fl/fl^* and *Mpc1^iCKO^* mice (Extended Data Fig. 1b-e). We anticipated that the contribution of glucose to TCA cycle metabolites and thus cardiac ATP production would decrease in *Mpc1^iCKO^* mice. Unexpectedly, we observed that the glucose carbon contribution to TCA cycle metabolites actually increased in *Mpc1^iCKO^*animals compared to controls (Fig. 1c). ^13^C labeling was highest in the M+1 isotopomers of succinate and malate, and present in M+2, indicating rapid TCA cycle turning (Extended Data Fig. 1f,g)^18^. The contribution of lactate carbon to TCA cycle metabolites was similar in *Mpc1^fl/fl^*and *Mpc1^iCKO^* hearts (Fig 1d. and Extended Data Fig. 1h,i). We performed additional infusions of other major cardiac fuels (palmitate, oleate and 3-hydroxybutyrate) to determine whether *Mpc1^iCKO^* mice displayed systemic alterations in cardiac fuel choice that may have confounded our glucose and lactate tracing results (Extended Data Fig. 1j-l). However, the nutrient-specific contributions to the cardiac TCA cycle in young *Mpc1^iCKO^* mice were not statistically different from controls (Extended Data Fig. 1j-m), suggesting that heart metabolism is resilient to loss of the MPC.

### Lactate is the preferred carbohydrate fuel of cardiomyocytes

We reasoned that direct import of lactate into mitochondria could explain the MPC-independence of carbohydrate oxidation in *Mpc1^iCKO^*animals. To understand how lactate is metabolized in the myocardium, but without the added complication of other cell types or the circulation, we turned to cultured primary adult cardiomyocytes (ACMs) (Extended Data Fig. 2a). Control ACMs were competent to oxidize both lactate and glucose as assessed by ^13^C incorporation into TCA cycle metabolites (Extended Data Fig. 2b). However, when unlabeled lactate was added to standard lactate-free ACM culture media containing [U-^13^C]glucose (DMEM), labeling from glucose was almost eliminated, suggesting a strong preference for lactate over glucose oxidation (Fig. 1e and Extended Data Fig. 2c,d). ACMs from *Mpc1^iCKO^* and *Mpc1^fl/fl^* mice metabolized lactate into TCA metabolites equivalently (Extended Data Fig. 2e). We tested whether the ACM preference for lactate would translate into increased cell fitness by incubating cultures in standard lactate-free DMEM or a lactate-containing Human Plasma-Like Media supplemented with physiological levels of BSA-conjugated fatty acids and calcium (HPLM-FA)^19^ (Extended Data Fig. 2f). We found that ACMs cultured in HPLM-FA had a substantial increase in average survival, from 2 to 7 days, and this was attenuated when lactate was removed (Fig. 1f). ACMs cultured in HPLM-FA retained inducible contractile function in culture for as long as 7 days (Supplemental Video 1).

MPC-independent mitochondrial oxidation requires a transporter to import lactate across the inner mitochondrial membrane (IMM). Prior reports suggested that the plasma membrane lactate transporter MCT1, encoded by *Slc16a1*, could be identified within mitochondria from cardiomyocytes^20,21^, however the concept of mitochondrial lactate metabolism is considered by many to be highly controversial^22,23^. We observed MCT1 protein by immunoblot in the mitochondrial fraction of *Mpc1^fl/fl^* hearts, and this was increased in *Mpc1^iCKO^*hearts (Fig. 1g,h, Extended Data Fig. 2g,h). To test whether MCT1 is necessary for lactate oxidation in mitochondria, we repeated our [U-^13^C]glucose tracing in *Mpc1^iCKO^* animals treated with a potent MCT1 inhibitor, AZD3965^24^. AZD treatment blocked the oxidation of glucose carbon as measured by TCA cycle metabolite labeling only in *Mpc1^iCKO^* and not control animals (Fig. 1i, Extended Data Fig. 1n-r). This result suggests the presence of functional MCT1 in mitochondria and demonstrates the redundancy of MCT1 and the MPC for the oxidation of glucose carbon in heart.

### MCT1 localizes to the IMM in ACMs and myocardial tissue

To explore the putative role of MCT1 as a mitochondrial lactate transporter, we developed a tamoxifen-inducible, cardiac-specific *MCT1* knockout mouse: *Mct1^fl/fl^*:αMHC^MerCreMer*+/-*^, hereafter *Mct1^iCKO^*. We validated knockout by PCR and MCT1 immunoblots from whole heart and mitochondrial preparations (Extended Data Fig. 3a,b). We isolated ACMs from the myocardium and performed immunofluorescence staining with MCT1 antibodies to determine subcellular localization. We observed distributed punctate staining throughout cardiomyocytes that co-localized with Mitotracker Red (Fig. 2a,b). MCT1 signal was highest at the edges of mitochondria and absent in *Mct1^iCKO^*ACMs. MCT1 staining co-localized with IMM protein SLC25A6, but not mitochondrial outer membrane marker TOMM20 (Fig. 2c,d). To independently validate MCT1 mitochondrial localization, we performed a proteinase K protection assay and found that MCT1 displayed the same distribution pattern as ATP5A, an integral IMM protein (Fig. 2e). We also identified lactate dehydrogenase LDHB, but not LDHA within the mitochondrial matrix (Fig. 2e, Extended Data Fig. 3b). Finally, we identified MCT1 in purified human cardiac mitochondria by MCT1 immunoprecipitation (IP) followed by immunoblot (Fig. 2f) and LC-MS (Extended Data Fig. 3c-f). Analysis of previously published single-nuclei RNAseq data showed that *SLC16A1* transcripts are predominantly expressed in cardiomyocytes from human hearts (Extended Data Fig. 3g)^25^.

**Fig. 2.**
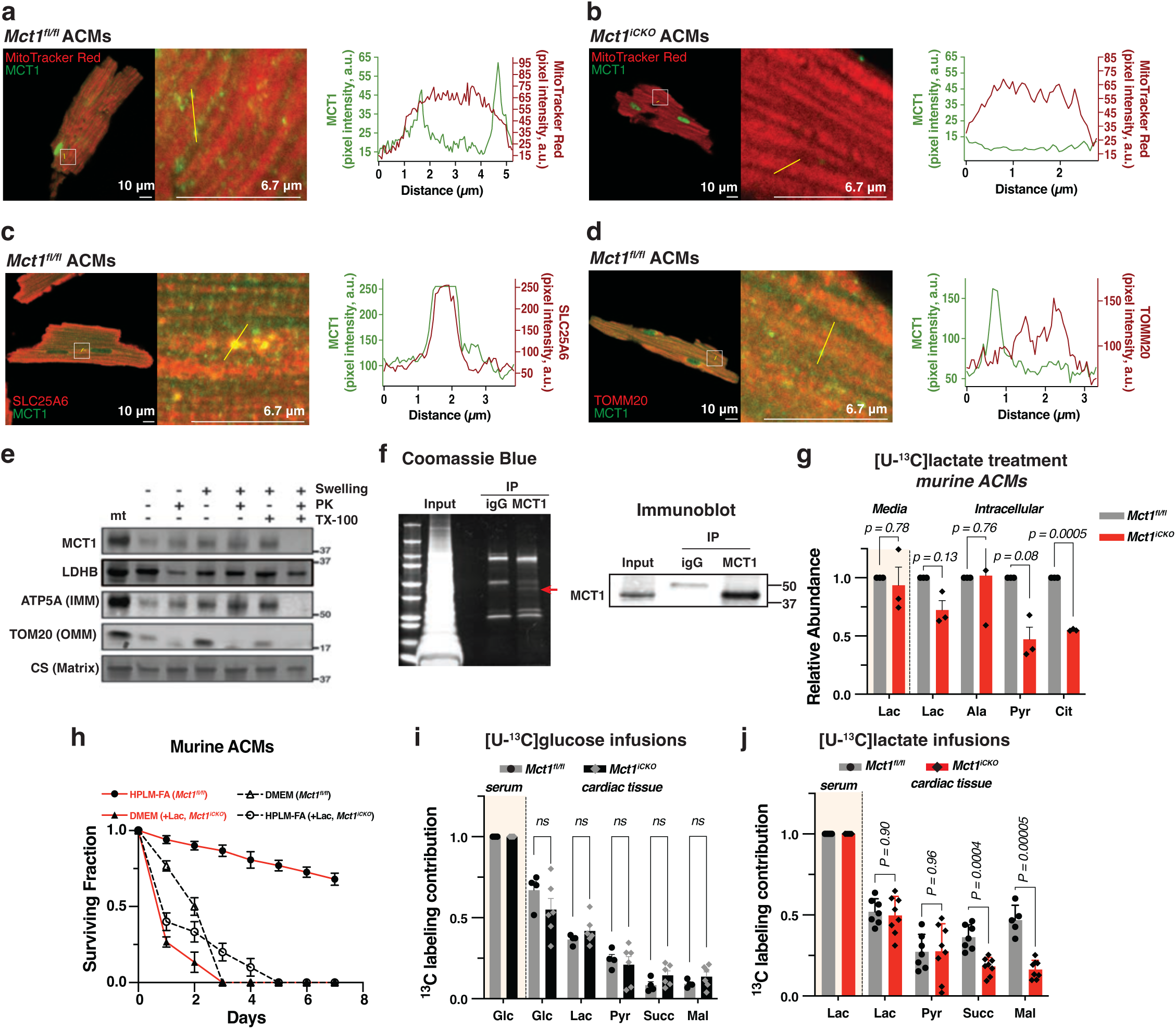
Lactate oxidation requires MCT1. **a**, **b**, Representative images of ACMs from *Mct1^fl/fl^*and *Mct1^iCKO^* mice labeled with MitoTracker Red CMXRos and co-stained for endogenous MCT1 (green). MCT1 and MitoTracker Red pixel intensity plots for the yellow line are graphed on the right of each panel. Scale bars represent 10 μM (full image) and 6.7 μM (magnified). **c**, Representative images of *Mct1^fl/fl^* ACMs stained for endogenous SLC25A6 (red) and MCT1 (green) and **d**, Endogenous TOMM20 (red) and MCT1 (green). Corresponding pixel intensity plots for the yellow line shown in the magnification are graphed on the right of each panel. Scale bars represent 10 μM (full image) and 6.7 μM (magnified). **e,** Immunoblot of Proteinase K protection assay conducted on mitochondria isolated from human cardiac tissue. **f**, Coomassie stained SDS-Page gel (left) of mitochondrial preparations isolated from human cardiac tissue immunoprecipitated for MCT1 or IgG and corresponding anti-MCT1 immunoblot (right). **g**, ^13^C enrichment in lactate, alanine, pyruvate or citrate from *Mct1^fl/fl^* or *Mct1^iCKO^* ACMs cultured with [U-^13^C]lactate. *n=3* independent ACM preparations from unique hearts. **h**, Survival fraction of *Mct1^fl/fl^* or *Mct1^iCKO^* ACMs grown in DMEM or HPLM-FA medias supplemented with lactate. **i**, Heart ^13^C labeling of glucose, pyruvate and TCA cycle metabolites from [U-^13^C]glucose infusions in *Mct1^fl/fl^*(*n=4*), or *Mct1^iCKO^*(*n=6*) mice. **j**, Heart ^13^C labeling of lactate and TCA cycle metabolites from [U-^13^C]lactate infusions in *Mct1^fl/fl^*(*n=7*) or *Mct1^iCKO^* (*n=8*) mice. Glc: glucose; Lac: lactate; Pyr: pyruvate; Succ: succinate; Mal: Malate. All data represent mean ± SEM. Significance determined by two-way ANOVA (g), and by multiple comparisons corrected (Holm-Sídák method) unpaired t tests (I,j). ns, not significant (p>0.05).

### MCT1 is required for the oxidation of lactate in ACMs and heart

To understand whether MCT1 has a functional role in lactate oxidation, we incubated ACMs from *Mct1^fl/fl^* (control) and *Mct1^iCKO^* animals with [U-^13^C]lactate. Lactate contributed to citrate (M+2) labeling in control ACMs, but this was sharply reduced in *Mct1^iCKO^* ACMs (Fig. 2g). Lactate M+3 labeling was not statistically different between genotypes indicating that altered citrate labeling was likely due to changes in oxidation and not substrate import in the cytoplasm (Fig. 2g, Extended Data Fig. 4a,b). Consistent with our hypothesis that lactate oxidation is mediated by MCT1, the viability of ACMs derived from *Mct1^iCKO^* hearts was not improved when cultured in lactate-containing media (HPLM-FA or DMEM) (Fig. 2h). To determine whether MCT1 was necessary for the oxidation of lactate *in vivo*, we infused *Mct1^iCKO^* animals with [U-^13^C]glucose or [U-^13^C]lactate. In *Mct1^iCKO^* hearts, the oxidation of lactate into TCA metabolites was significantly reduced, whereas labeling from glucose was unchanged (Fig. 2i,j, Extended Data Fig 4c-h). Serum lactate and glucose metabolic parameters were not affected (Extended Data Fig. 4i-l). Importantly, loss of MCT1 did not alter tissue lactate or pyruvate enrichment, suggesting that the observed TCA cycle defects were not due to a loss of cellular lactate import. Together, these data show that cardiac lactate oxidation is dependent upon cardiomyocyte MCT1.

### MCT1 is required for purified mitochondria to respire on lactate

We purified mitochondria from *Mct1^fl/fl^* (control), *Mct1^iCKO^* and *Mpc1^iCKO^* hearts and characterized their respiration and metabolism on different substrates. Mitochondria from control and *Mct1^iCKO^* hearts respired equivalently on pyruvate, but *Mpc1^iCKO^* mitochondria had reduced oxygen consumption rates (*JO_2_*), impaired ^13^C-citrate production and suppressed pyruvate entry (Fig. 3a,b, Extended Data Fig. 5a). Conversely, respiration on lactate was similar in both control and *Mpc1^iCKO^* mitochondria but was reduced in mitochondria from *Mct1^iCKO^* hearts (Fig. 3c). [U-^13^C]lactate import and oxidation into TCA cycle metabolites was attenuated in *Mct1^iCKO^*mitochondria (Fig. 3d, Extended Data Fig. 5b-e). In contrast, we observed normal lactate metabolism in mitochondria isolated from the hearts of mice lacking the cardiomyocyte lactate exporter MCT4 (Extended Data Fig. 5f-j)^17^. Mitochondria from *Mct1^iCKO^* mice showed no defect in mitochondrial electron transport chain complex or supercomplex assembly (Extended Data Fig. 5k). Acute treatment of mitochondria isolated from control animals with an MCT1 inhibitor (7ACC2) or a pan-LDH inhibitor (GSK 2837808A) inhibited respiration on lactate (Fig. 3e). Finally, we treated purified mitochondria from human non-failing donor hearts with [U-^13^C]lactate and observed impaired oxidation to pyruvate and citrate when treated with an MCT1 inhibitor but not with an MPC inhibitor (Fig. 3f). We noted that total oxygen consumption on lactate was reduced compared to pyruvate, suggesting incomplete lactate oxidation. We hypothesized that this was due in part to mitochondrial pyruvate efflux via the MPC that was occurring in our buffers lacking pyruvate. Indeed, administrating the MPC inhibitor UK5099 to ACMs prior to lactate addition enhanced mitochondrial lactate respiration (Extended Data Fig. 5l-o).

**Fig. 3.**
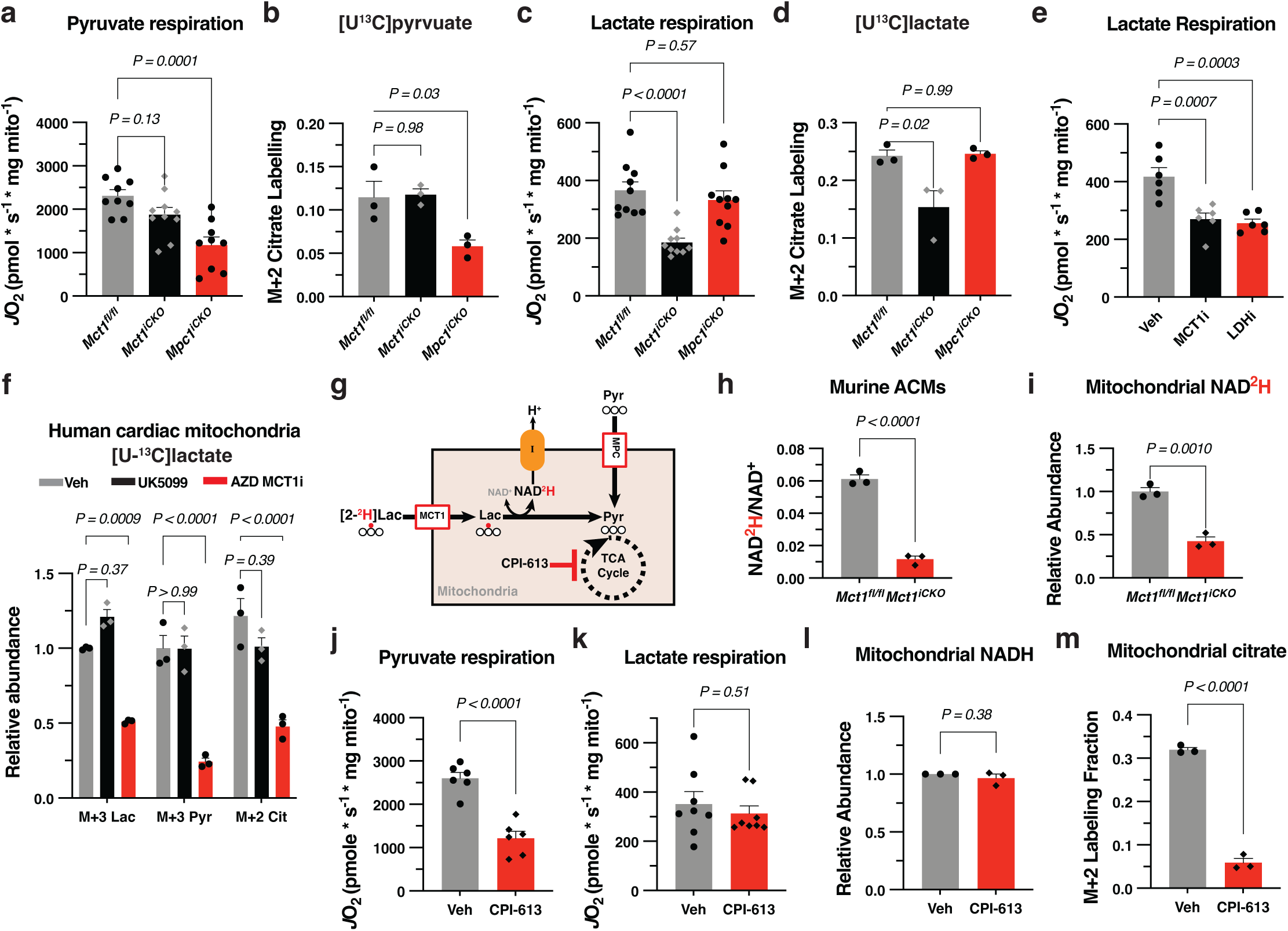
Mitochondrial MCT1 is necessary for respiration on lactate. **a**, Oxygen consumption rates (*J*O_2_) of isolated cardiac mitochondria incubated with pyruvate from the indicated genotypes (*Mct1^fl/fl^ n=9*, *Mct1^iCKO^ n=10*, *Mpc1^iCKO^ n=9*). **b**, Citrate M+2 labeling fraction from isolated cardiac mitochondria incubated with [U-^13^C]pyruvate (*n=3* individual hearts). **c**, Oxygen consumption rates of isolated cardiac mitochondria incubated with lactate (WT *n=10*, *Mct1^iCKO^ n=10*, *Mpc1^iCKO^ n=10*). **d**, Citrate M+2 labeling fraction from isolated cardiac mitochondria incubated with [U-^13^C]lactate (*n=3* individual hearts). **e**, Oxygen consumption rates of isolated cardiac mitochondria incubated with lactate treated with Veh (DMSO) *n=6*, MCT1i (7ACC2) *n=6*, LDHi (GSK 2837808A) *n=6*. **f**, ^13^C labeling fractions of lactate (M+3), pyruvate (M+3) and citrate (M+2) from human donor cardiac mitochondria incubated with [U-^13^C]lactate (*n*=*3* human hearts). **g**, Schematic illustrating the transfer of electrons from lactate (traced by ^2^H hydride transfer) into the mitochondrial NADH pool for respiration. **h**, Ratio of NADH (M+1) over NAD^+^ from ACMs incubated with [2-^2^H]lactate (*n=3* for *Mct1^fl/fl^* and *Mct1^iCKO^*). **i**, Normalized NADH (M+1) ion intensities from isolated mitochondria incubated with [2-^2^H]lactate (*n=3* for *Mct1^fl/fl^* and *Mct1^iCKO^*). **j, k,** Oxygen consumption rates of isolated cardiac mitochondria incubated with pyruvate or lactate and treated with DMSO (Veh), or CPI-613 (*n=6* hearts for pyruvate, *n=8* hearts for lactate). **l**, Normalized NADH ion intensities from isolated mitochondria treated with DMSO (Veh), or CPI-613, *n=3*. **m**, Citrate M+2 labeling fraction from isolated cardiac mitochondria incubated with [U-^13^C]lactate and treated with DMSO (Veh) or CPI-613, *n=3*. All data represent mean ± SEM. Significance determined by one-way ANOVA with Dunnett’s multiple comparison test (a-e), two-way ANOVA (f) and by unpaired two-tailed t-tests (h-m). ns, not significant (p>0.05).

### Direct lactate import supports TCA cycle-independent respiration

The oxidation of one lactate molecule within the mitochondria by LDH to produce pyruvate generates one molecule of NADH that can transfer electrons directly into the electron transport chain (Fig. 3g). To test whether LDH activity is sufficient to sustain respiration even in the absence of TCA cycle metabolism, we incubated control and *Mct1^iCKO^* ACMs with [2-^2^H]lactate and observed MCT1-dependent ^2^H labeling of NADH (Fig. 3h). We observed a similar MCT1 dependent transfer of ^2^H from [2-^2^H]lactate to NADH in mitochondria isolated from these mice (Fig. 3i). To test whether this LDH activity could generate sufficient NADH for respiration, we inhibited the TCA cycle with the pyruvate- and α-ketoglutarate-dehydrogenase inhibitor CPI-613^26^. As expected, mitochondrial respiration on pyruvate was sharply reduced by CPI-613 treatment, however lactate respiration was unaltered (Fig. 3j-l). This occurred even as M+2 citrate production from ^13^C-lactate was suppressed (Fig. 3m). Collectively, these data suggest the presence of LDH activity within the matrix sufficient to power mitochondrial respiration on lactate.

### MCT1 loss accelerates the development of heart failure

Having established that MCT1 mediates mitochondrial lactate import and oxidation, we asked whether loss of this biochemical functionality impacts cardiac physiology. *Mct1^iCKO^* mice did not show evidence of structural or functional cardiac abnormalities by 1 year of age as measured by serial echocardiography (Extended Data Fig. 6a-i). Consistent with this, *Mct1^iCKO^*mice exhibited unchanged maximal exercise capacity over time (Extended Data Fig. 6j,k). We hypothesized that the absence of a phenotype was a consequence of the functional redundancy between MCT1 and MPC in cardiomyocytes. Therefore, we turned to neurohormonal agonist-induced models of cardiac hypertrophy that we previously reported to decrease *Mpc1* expression^17^. We surgically implanted mice with osmotic minipumps delivering angiotensin II and phenylephrine (Ang/PE) to induce cardiac hypertrophy. After 6 weeks, all treated animals developed cardiac hypertrophy, but it was more pronounced in *Mct1^iCKO^*mice (Fig. 4a and Extended Data Fig 7a). As we anticipated, Ang/PE treatment led to a large reduction in *Mpc1* expression, and this was observed in both *Mct1^iCKO^*and *Mct1^fl/fl^* animals (Fig. 4b). Body weight, heart rate (HR), left ventricular end systolic diameter (D;s) and left ventricular end diastolic diameter (LVEDD) were not significantly different between genotypes (Extended Data Fig. 7b-e). In contrast, systolic function as measured by stroke volume (SV) was significantly reduced in *Mct1^iCKO^* animals leading to reduced cardiac output (CO), fractional shortening (FS) and impaired left ventricular ejection fraction (LVEF) (Fig. 4c and Extended Data Fig. 7f-i). Consequently, Ang/PE treated animals showed reduced overall survival which did not reach statistical significance likely due to the relatively short follow-up period (Extended Data Fig. 7j). Similarly, loss of MCT1 accelerated the development of heart failure (HF) in mice when cardiac hypertrophy was induced by transverse aortic constriction (TAC) (Fig. 4d and Extended Data Fig. 8a-k.).

**Fig. 4.**
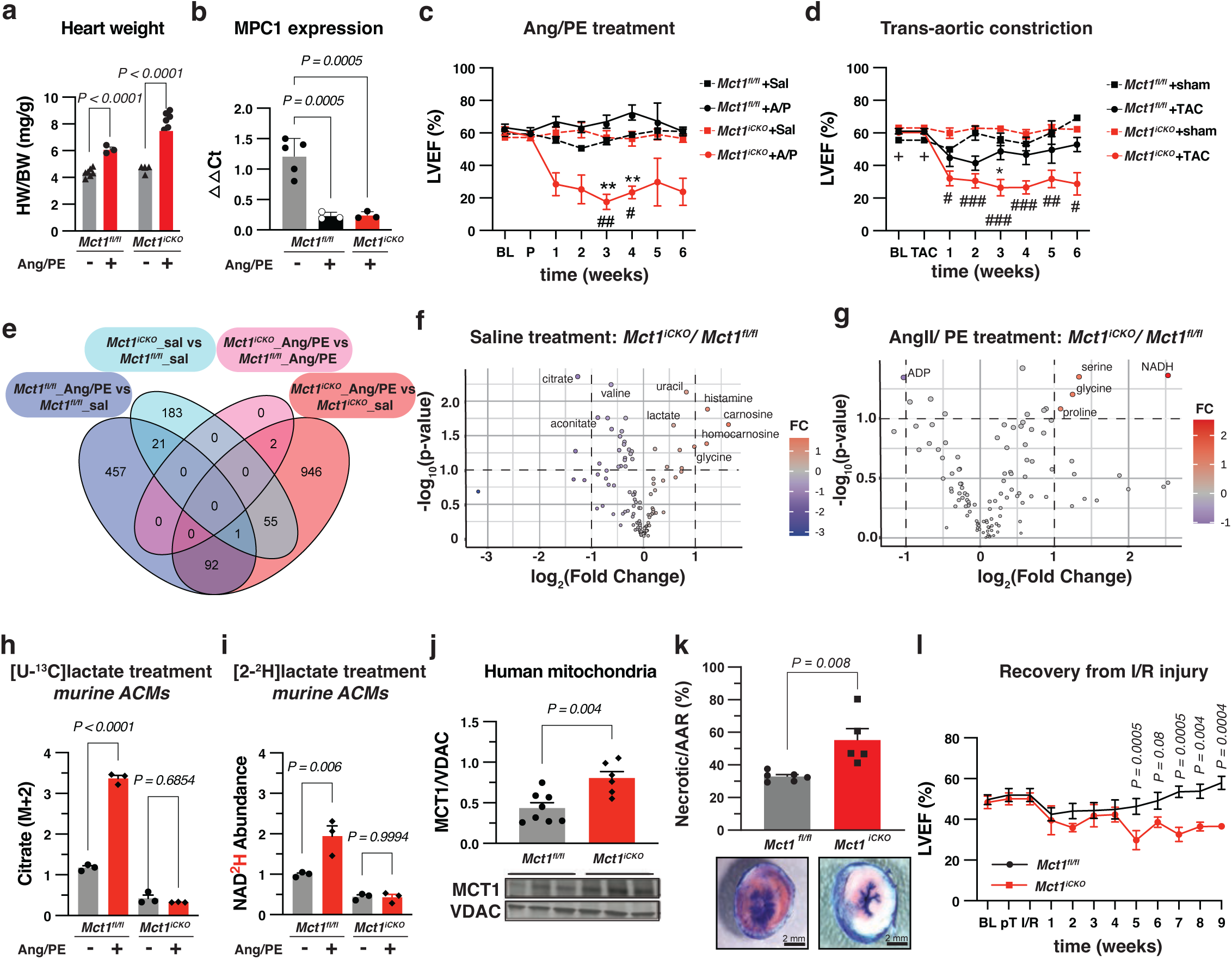
Loss of MCT1 impairs cardiac function upon injury. **a**, Heart weight to body weight ratio of mice treated with angiotensin II and phenylephrine (Ang/PE) by osmotic minipump for 42 days (*Mct1^fl/fl^* saline *n*=7, *Mct1^fl/fl^* Ang/PE *n*=3, *Mct1^iCKO^* saline *n*=4, *Mct1^iCKO^* Ang/PE *n*=6). **b**, Quantification of *Mpc1* transcript levels from Ang/PE treated hearts (*Mct1^fl/fl^* saline *n*=5, *Mct1^fl/fl^*Ang/PE *n*= 3, *Mct1^iCKO^* Ang/PE *n*=3). **c**, Left ventricular ejection fraction (LVEF) of same Ang/PE mice (*Mct1^fl/fl^*saline *n=7*, *Mct1^fl/fl^* Ang/PE *n=8*, *Mct1^iCKO^*saline *n=4*, *Mct1^iCKO^* Ang/PE *n=10*). Statistical significance was determined by a mixed-effects model (repeated measures ANOVA) with the Geisser-Greenhouse correction was used. If significant, then a Tukey’s multiple comparison test was applied with individual variances computed for each comparison. *=p<0.05, **=p<0.01 comparing *Mct1^fl/fl^*Ang/PE vs. *Mct1^iCKO^* Ang/PE. #=p<0.05, ##=p<0.01, comparing *Mct1^iCKO^* Ang/PE vs. *Mct1^iCKO^* saline. **d**, LVEF of mice subjected to trans-aortic constriction (TAC) and monitored for 6 weeks (*Mct1^fl/fl^* sham *n*=5, *Mct1^fl/fl^* TAC *n*=9, *Mct1^iCKO^* sham *n*=4, *Mct1^iCKO^* TAC *n*=10). Statistical significance was determined by a mixed-effects model as in c. *=p<0.05, **=p<0.01 comparing *Mct1^fl/fl^*TAC vs. *Mct1^iCKO^* TAC. #=p<0.05, ##=p<0.01, ###=p<0.001, comparing *Mct1^iCKO^* TAC vs. *Mct1^iCKO^* sham. +=p<0.05, comparing *Mct1^fl/fl^*sham vs. *Mct1^iCKO^* sham. **e**, Venn diagram showing intersection of differentially expressed genes between hearts from *Mct1^iCKO^* and *Mct1^fl/fl^* Ang/PE and vehicle treated animals (*Mct1^fl/fl^* saline *n=6*, *Mct1^fl/fl^* Ang/PE *n=3*, *Mct1^iCKO^* saline *n=4*, *Mct1^iCKO^* Ang/PE *n=5*). **f**, Volcano plot showing differential abundance of polar metabolites in saline treated *Mct1^iCKO^* and *Mct1^fl/fl^* hearts (*Mct1^fl/fl^* saline *n=7*, *Mct1^iCKO^*saline *n*=4). **g**, Volcano plot showing differentially abundant polar metabolites in Ang/PE treated *Mct1^iCKO^* and *Mct1^fl/fl^* hearts (*Mct1^fl/fl^* Ang/PE n=3, *Mct1^iCKO^* Ang/PE *n=5*). **h**, M+2 citrate labeling fraction from ACMs cultured with [U-^13^C]lactate and treated with Ang/PE (*n=3*). **i**, Normalized NADH (M+1) ion intensities from ACMs cultured with [2-^2^H]lactate, n=3. **j**, Quantified MCT1 immunoblot bands from mitochondria isolated from human cardiac tissue (donor *n=8*, HF *n=6*). **k**, Evans Blue staining of heart sections showing myocardial salvage and necrosis following in vivo I/R injury and quantification of necrotic tissue within the area at risk (*n=5*). **l**, LVEF of mice subjected I/R injury and their recovery over 9 weeks (*Mct1^fl/fl^ n=11*, *Mct1^iCKO^ n=9*). Data are plotted as mean ± SEM. Significance determined by unpaired two-tailed t-test (j,k) with multiple comparisons correction (l) or one-way ANOVA with Dunnett’s multiple comparison test (a,b, and h-i). ns, not significant (p>0.05).

To understand the accelerated development of HF in *Mct1^iCKO^*animals, we performed transcriptomic and metabolomic analyses on Ang/PE-treated hearts. Few genes were significantly different between *Mct1^iCKO^* and control animals at baseline but Ang/PE treatment had a large effect on the transcriptome in both genotypes (Fig. 4e and Extended Data Fig. 9a-c). However, Ingenuity Pathway Analysis revealed genotype specific differences in pathway activation upon Ang/PE treatment with “*contractility of cardiac muscle*” most impacted in *Mct1^iCKO^*mice (Extended Data Fig. 9d,e). Surprisingly, when comparing Ang/PE treated *Mct1^iCKO^* and *Mct1^fl/fl^* hearts, only 2 significantly dysregulated genes were observed suggesting a role for non-transcriptional mechanisms in cardiac dysfunction (Fig. 4e). Indeed, metabolomic analysis of the same hearts revealed significant genotype-specific differences between *Mct1^iCKO^* and *Mct1^fl/fl^* hearts both with saline and Ang/PE treatment (Fig. 4f,g and Extended Data Fig. 9f-j). *Mct1^iCKO^* hearts had elevated lactate, histamine, carnosine, glycine and reduced levels of citrate/isocitrate compared to *Mct1^fl/fl^* hearts with saline treatment. Upon Ang/PE treatment, NADH was the most upregulated metabolite in *Mct1^iCKO^* hearts, suggestive of impaired oxidation in these animals (Fig. 4g). Consistent with a critical mitochondrial defect, all purine nucleotides were downregulated, with ADP being the most statistically significantly reduced metabolite in failing *Mct1^iCKO^* hearts (Fig. 4g). Metabolome differences were much less pronounced upon Ang/PE treatment in control hearts (Extended Data Fig. 9k).

### Cardiomyocyte lactate metabolism is increased by neurohormonal agonists

Our transcriptomic and metabolic profiling of Ang/PE-treated *Mct1^iCKO^*hearts was completed only after the development of significant cardiac dysfunction. To ask what the acute effects of neurohormonal stimulation were upon metabolism, we treated *Mct1^fl/fl^* and *MCT1^iCKO^*ACMs for 48 hours with AngII/PE and observed a robust decrease in viability and increase in cell size in both genotypes (Extended Data Fig. 10a-c). We next co-cultured treated cells with [U-^13^C]lactate and observed an increase in citrate M+2 labeling fraction that was dependent upon MCT1 expression (Fig. 4h). Similarly, Ang/PE treatment increased the [2-^2^H]lactate contribution to ^2^H NADH in *Mct1^fl/fl^* but not *Mct1^iCKO^* ACMs (Fig. 4i). Reinforcing the importance of mitochondrial MCT1 in the metabolism of the failing heart, analysis of mitochondria from patients with HF with reduced ejection fraction and non-failing donor controls revealed increased mitochondrial MCT1 without a statistically significant increase in either total protein or transcript levels (Fig. 4j, Extended Data Fig. 10d,e).

### Loss of MCT1 worsens acute cardiac injury and potentiates the development of HF

Prior studies reported that MCT1 expression increases in myocardium during ischemia/ reperfusion (I/R) injury^27,28^. To study the consequences of MCT1 loss during I/R injury and to examine whether MCT1 has a role in oxygen-limited conditions, we studied *Mct1^iCKO^*mice subjected to I/R^29^ (Extended Data Fig. 10f). *Mct1^iCKO^* mice had myocardial areas at risk for infarction of similar size, but larger necrotic areas and a trend towards higher mortality compared to control littermates (Fig. 4k and Extended Data Fig. 10g-i). To understand whether these changes were correlated with intrinsic defects in *Mct1^iCKO^* cardiomyocyte function upon hypoxia we utilized an *in vitro* hypoxia-reoxygenation setup that models I/R injury^19^ (Extended Data Fig. 10j). At baseline, *Mct1^iCKO^* ACMs had increased mitochondrial membrane potential (ΔΨ), higher calcium (Ca^2+^) levels, and elevated reactive oxygen species (ROS) that were further increased after hypoxia-reoxygenation (Extended Data Fig. 10k-m). The significant increase in ΔΨ and ROS suggests a heightened energetic state in *Mct1^iCKO^* ACMs which may exacerbate cellular damage and contribute to poorer outcomes during I/R injury.

Collectively, our data indicated a key role for cardiac MCT1 in metabolic adaptation to models of both acute and chronic cardiac stress. To assess whether MCT1 was important for recovery from injury and progression of cardiac remodeling, we allowed *Mct1^iCKO^* mice subjected to I/R injury to recover (Extended Data Fig. 11a). In both *Mct1^fl/fl^* and *Mct1^iCKO^*mice, LVEF fell immediately after the procedure and by the same amount despite the larger injury in the knockout animals (Fig. 4l). However, in *Mct1^fl/fl^* mice systolic function returned to baseline by 7 weeks whereas *Mct1^iCKO^* mice showed no recovery of heart function for the duration of the study (Fig. 4m and Extended Data Fig. 11b-d).

## Discussion

The data presented here demonstrate that in the myocardium glucose carbons can enter the TCA cycle independently of the MPC, and this depends on the lactate transporter MCT1. Using a combination of mouse genetic models and inhibitors, we show that MCT1 resides in the IMM and its loss leads to functional defects in mitochondrial, cellular and tissue oxidation of lactate. Loss of MCT1 impairs metabolic adaptation to cardiac stress and leads to significant cardiac structural and functional impairment and HF. Based on these data, we propose that mitochondrial import of lactate serves to facilitate lactate carbon feeding the TCA cycle without disrupting the cytosolic redox balance and impacting glucose metabolism. Our model unifies recent results that demonstrate high lactate oxidation fluxes in mammalian physiology^6,7,30^ with results from cellular lactate biosensors showing high concentrations of lactate within mitochondria^31,32^.

Our data are broadly consistent with the ‘lactate shuttle’ hypothesis developed by physiologists and supported by data from both organismal and cell systems^33–37^. By positioning the LDH reaction within mitochondria, lactate can shuttle an additional pair of cytosolic electrons into mitochondrial NADH for transfer into the mitochondrial quinone (Q) pool to support complex I activity (Fig 3h,i). When coupled with MPC-mediated pyruvate export, this MCT1-dependent electron shuttle can sustain respiration even in the absence of a functional TCA cycle (Fig 3j,k). ATP production from lactate depends upon oxygen to serve as the terminal electron acceptor and drive the equilibrium of the reaction towards carbon oxidation. The lactate shuttle hypothesis, however, has been vigorously contested based on conflicting data from isolated mitochondrial systems^31^ and arguments about the equilibrium thermodynamics of the LDH reaction^22,23^. By presenting the first fully profiled genetic model to test the lactate shuttle hypothesis, our work demonstrates that mitochondrial lactate oxidation best supports functional and tracing data from purified mitochondria, cells and organs.

The rapid uptake and oxidation of lactate has been well-documented in exercise physiology, underscoring lactate’s vital role as a supply of systemic energy under aerobic exercise demand^38–40^. However, lactate oxidation to pyruvate is not compatible with high glycolytic flux, requiring the invocation of compartmentalized metabolism^8,22^. Placing lactate oxidation within the mitochondria of the myocardium solves this conundrum, leading to greater adaptability and taking advantage of the expression of LDHB (heart isoform) which has a reduced *K_M_* for pyruvate^41^, and is thus better suited to lactate oxidation compared to LDHA. Mitochondrial lactate transport enables not only simultaneous lactate and glucose oxidation, but also provides crucial redundancy for carbohydrate fuel entry into the mitochondria, enabling cardiac cells to maintain glucose oxidation even in the absence of the MPC^42,43^. We show that this redundancy is critical to support the heart’s need for a rapid and flexible energy supply in the face of cardiac stress. In conclusion, our findings underscore the significance of mitochondrial lactate metabolism to cardiac structure and function, adaptation and survival.

## Methods

### Human heart samples

Human myocardial tissue samples were acquired from patients through the Utah Cardiac Recovery Program at the University of Utah under an approved IRB protocol (IRB-00030622). All samples were collected with informed consent from patients, organ donors, or their guardians. Samples were acquired from consented patients who had chronic advanced heart failure with reduced ejection fraction at the time of heart transplantation. Non-failing donors that were ineligible for heart transplant due to non-cardiac reasons (donor-recipient size mismatch, infection, etc.) were used as a control. Myocardial tissue was collected in the operating room and immediately frozen in liquid nitrogen and subsequently stored in −80°C until further use.

### Animal care

All procedures were approved by the Institutional Animal Care and Use Committee (IACUC) of the University of Utah. Mice were housed at 22–23°C using a 12 hour (hr) light/12 hr dark cycle. Animals were maintained on irradiated chow diet (Teklad Diet 2920x) with ad libitum access to water at all times. Food was only withdrawn during experiments starting at the beginning of the light cycle.

Adult cardiac-specific MPC1-knock out mice were generated as previously described^44^. *Mct1^fl/fl^*were generated as previously described^45^. Cardiac specific *Cre*-recombinase was introduced by crossing mice with αMHC^MerCreMer^ animals (Jackson Labs strain #005657). 8-week-old *Mct1^fl/fl^*:αMHC^MerCreMer*+/-*^ and *Mct1^fl/fl^*:αMHC^MerCreMer*-/-*^ littermates were injected intraperitoneally with 40 mg/ kg of body weight tamoxifen for three consecutive days to yield adult *Mct1^iCKO^* and *Mct1^fl/fl^*(control) littermates.

Adult cardiac-specific MCT4-knock out mice were generated as follows. Three embryonic clones carrying *Slc16a3 tm1a* (solute carrier family 16, monocarboxylic acid transporter member 3 known as MCT4) were obtained from EUMMCR (the European Conditional Mouse Mutagenesis repository, http://www.eummcr.org/). All clones were from the cell line JM8A3.N1 (background strain C57Bl/6NTac with agouti coat color distribution). These ES clones were injected into the blastocoel of 3.5day old mouse blastocysts host strain C57Bl/6 Tyr c-Brd). Injected embryos were transferred to the uterine horns of appropriately timed pseudo pregnant recipient C57Bl/6 Tyr c-Brd females. The chimeras with highest ES contribution were bred with C57Bl/6J mates to test for germline transmission of the targeted allele. The pups were screened for the presence of germline transmitted tm1a and those positive were further propagated (first allele knockout tm1a, reporter-tagged insertion with conditional potential). Animals carrying the appropriate flox alleles were crossed with αMHC^MerCreMer^ ^mice^ as described above. Experiments on all animals began 4 weeks post-injection (12 weeks old).

### Echocardiographic assessment

Mice were anesthetized with 1.5% Isoflurane (Vet One, NDC 13985-046-60). Echocardiographic images were captured using the Vivo system sequentially. Both 2D long-axis and short-axis views were obtained and analyzed with Vivo Strain software (version 3.1.1). All measurements were conducted using two consecutive cardiac cycles. Limb leads were affixed using conductive cream to facilitate electrocardiogram (ECG) recording.

### Osmotic minipump model

12-week-old C57BL/6 mice were divided into 4 groups, weighed, and screened before surgery by echocardiography to establish baseline cardiac parameters. Surgical instruments were sterilized using a glass bead sterilizer prior to the surgery and aseptic technique was utilized during the procedure. The osmotic minipumps (Alzet, model 2006) were filled with Angiotensin II (1.5 μg/g/day) and Phenylephrine (50 μg/g/day) in 0.9% NaCl (0.15μl/hour for days) according to manufacturer instructions. The pumps were surgically implanted subcutaneously on the back through a small incision, under isoflurane (1.5% in O_2_) anesthesia. After the pump was implanted, and the incision was closed, the animals were placed in a clean, warm cage and monitored for any signs of distress. Next, mice were weighed and screened weekly via echocardiography. After 42 days of treatment with AngII/PE, the mice were sacrificed by cervical dislocation and blood and heart tissue were taken for further analysis.

### Trans-Aortic Constriction (TAC) experiments

Baseline echocardiograms were acquired, and transverse aortic constriction (TAC) was surgically performed on both male and female *Mct1^fl/fl^*and *Mct1^iCKO^* mice aged 12–14 weeks, as previously published^46,47^. Briefly, before surgery, a subcutaneous dose of sustained-release Buprenorphine SR (0.15 mg/kg) was administered for analgesia. Mice were anesthetized with 2-3% isoflurane in oxygen, delivered via nosecone connected to a VetFlo vaporizer. The aortic arch was visualized and constricted using a titanium clip. The surgical incision was closed using 6-0 sutures in layers. During recovery, regular-release buprenorphine was given as needed, for up to 48 hrs. Unclipped *Mct1^fl/fl^* and *Mct1^iCKO^*mice (sham) served as controls, undergoing anesthesia and a small incision without aortic constriction or cardiac damage. Following surgery, animals received weekly echocardiograms for six consecutive weeks to assess cardiac function and structure.

### Ischemia-reperfusion experiments

The experimental protocol was conducted on 20-30 g, 12–14-week-old *Mct1^iCKO^* and *Mct1^fl/fl^* mice, four weeks after tamoxifen injection and before the onset of HF symptoms. Mice were anesthetized with 2-3% isoflurane, intubated using a 22 GIV catheter, and ventilated at 100 breaths/min with a small rodent respirator. Mice were placed supine on a warming pad at 38°C, and chest hair was removed with depilatory cream, which was rinsed off with water or saline to avoid skin irritation. The skin was then prepped with betadine and alcohol. A thoracotomy was performed to access the heart, and a retractor was used to improve visibility. The pericardium was carefully removed, avoiding myocardial damage. Under a dissecting microscope, the left anterior descending coronary artery (LAD) was located, and a 6-0 silk suture was placed under it. A loose double knot was tied with a 2-3 mm diameter loop, through which a 2-3 mm piece of PE-10 tubing was placed. The tubing, pre-soaked in 100% ethanol and rinsed with sterile water, was used to tighten the loop around the artery, achieving coronary occlusion for 30 minutes. Successful LAD ligation was confirmed by a paler color on the LV’s anterior wall. After 30 minutes, the knot was untied, and the tubing was removed, confirming reperfusion by the return of a red color to the LV wall, which continued for 2 hrs.

In acute experiments, the coronary artery was briefly re-occluded after injury, and Evans blue dye was injected into the left atrium to mark the area at risk (AAR). Hearts were excised, perfused with saline, weighed, and sectioned into 1 mm slices using a vibratome, or separated into ischemic and non-ischemic samples and flash-frozen in liquid nitrogen. Slices were incubated in 1% triphenyltetrazolium chloride (TTC) at 38°C for 20 minutes to stain viable myocardium, then fixed in 10% formalin to distinguish between viable and necrotic tissue. Tissue sections were imaged the next day, and myocardial salvage was quantified using ImageJ by blinded researchers. For chronic experiments, mice underwent 30 minutes of ischemia, after which the ligature was removed, and the incisions were closed with sutures. Mice were placed in a warm cage, monitored for distress, weighed, and screened weekly via echocardiography.

### Exercise capacity testing

12–14-week-old mice (*Mct1^fl/fl^* and *Mct1^iCKO^*) practiced a 5-day treadmill acclimation protocol prior to any maximal exertion. This allowed for familiarity with the procedure, but did not produce any training effect on the animals^48^. Briefly, following a day of rest, mice were weighed, and the maximal treadmill test was performed. The test started with a warm-up period of 10 minutes, a treadmill speed of 10 m/min, and an incline of 10% grade. After this, the speed was increased by 5 m/min every 2 minutes until the mouse resisted a physical stimulus (touching the tail/backside of the mouse with a bristle brush) to run for more than 20 seconds or sat on the back of the non-moving treadmill belt 5-times in a 30-s period. If either occurred the time was recorded, the test was finished, and the mouse was returned to its cage to recover. Maximal treadmill speed did not exceed 30 m/min throughout the exercise testing. Total work (kg/m) and time to exhaustion (min) were then calculated for each mouse. Blood lactate (mm/L) was measured at 10, 20, and 30 m/min, and 10% grade to assess the systemic effects of lactate production in the *Mct1^iCKO^* compared with the *Mct1^fl/fl^*controls.

### Jugular vein catheterization surgery

Sterile mouse jugular vein catheters were purchased from SAI Infusion Technologies and one channel vascular access button from Instech Laboratories, Inc. Mice were fasted before surgery, pre-operatively shaved and administered bupivacaine before anesthesia. A dissection microscope (Leica S9i Stereo) was utilized during the placement of the catheter. Mice were provided standard recovery support and allowed at least 5 days recovery before further use in experiments. Mice with impaired vital signs, activity or reflexes were euthanized and not included in experimental studies.

### Stable isotope infusions

12-week-old surgically catheterized mice were maintained on a normal light cycle (6 AM – 6 PM). On the day of infusion experiment, mice were transferred to new cages without food at 6 AM (beginning of their sleep cycle) and infused for 2-3 hrs (metabolite-specific) starting at around 12 PM to reach serum steady-state based on previously published studies. The infusion setup (Instech Laboratories) included a swivel and tether to allow the mouse to move around the cage freely. Water-soluble isotope-labeled metabolites (Cambridge Isotope Laboratories, Tewksbury, MA) were prepared as solutions in sterile normal saline. To make ^13^C-labeled fatty acid solutions, the fatty acids were complexed with bovine serum albumin in a molar ratio 4:1. The infusion rate was set to 0.1 μL min−1 g−1 for water-soluble metabolites and 0.4 μL min−1 g−1 for fatty acids. Final infusion solutions and times were 200 mM [U-^13^C]glucose for 3 hours, 490 mM [U-^13^C]sodium lactate for 2 hours, 50 mM [U-^13^C]3-hydroxybutyrate, 8 mM [U-^13^C]oleic acid or 4 mM [U-^13^C]sodium palmitate all for 3 hours. [U-^13^C]glucose infusion with AZD3965 treatment was achieved by dosing mice with 100 mg/kg AZD3965 or vehicle (0.5% HPMC, 0.1% Polysorbate 80) by oral gavage at 6 hrs prior to infusion start and again at start of infusion. [U-^13^C]glucose was infused using the same parameters as other experiments and mouse blood and tissue harvested as described. Blood was collected by tail snip (∼10 μL) and transferred into blood collection tubes (Microvette CB 300 Z). Blood samples were stored on ice and then centrifuged at 5,000 × g for 5 minutes at 4°C to collect serum. Tissue harvest was performed at the end of the infusion after euthanasia by cervical dislocation. Tissues were quickly dissected, rinsed in cold PBS, clamped with a pre-chilled Wollenberger clamp, and dropped in liquid nitrogen.

### Tissue and serum metabolite extraction

Serum (4 μL) was diluted with 140 μL of ice cold 80% methanol:20% water solution and vortexed. 72 μL of chloroform was added, the solution vortexed again and centrifuged at 15,060 × g for 10 minutes at 4°C to enforce phase separation. Aqueous supernatant was used for LC-MS analysis. For tissue extractions, 30-40 mg sections of snap frozen mouse tissue were transferred to pre-chilled Safe-Lock tubes (Eppendorf, 022363352) containing a cold 5/16 in. diameter stainless steel ball (Grainger, 4RJL8). The tissue was disrupted by shaking at 25 Hz for 30 sec under liquid nitrogen using the Retsch CryoMill (Retsch, 20.749.0001). 15 μL per mg of tissue of −80°C polar metabolite extraction solution containing 40:40:20 Acetonitrile:Methanol:Water and 0.1% Formic Acid was added to homogenized tissue. Samples were briefly vortexed before neutralizing with 8 μL of 15% ammonium bicarbonate per 100 mL of extraction solvent. The solution was centrifuged at 16,000 × g for 10 minutes at 4°C. The supernatant was collected and the pellet extracted a second time with 525 μL of chilled 40:40:20 methanol:acetonitrile:water. Both supernatants were combined and 525 μL of chloroform was added. The mix was vortexed and centrifuged at 16,000 × g for 10 minutes at 4°C, and final aqueous layer transferred to LC-MS tubes for analysis.

### LC-MS analysis of polar metabolites

Extracted polar metabolite samples were analyzed by LC-MS. Separation was achieved by hydrophilic interaction liquid chromatography (HILIC) using a Vanquish HPLC system (ThermoFisher Scientific). The column was an Xbridge BEH amide column (2.1 mm x 150 mm, 2.5 µm particle size, 130 Å pore size, Waters) run with a gradient of solvent A (20 mM ammonium hydroxide, 20 mM ammonium acetate in 95:5 acetonitrile:Water, pH 9.5) and solvent B (100% acetonitrile) at a constant flow rate of 150 μL/min. The gradient function was: 0 min, 90% B; 2 min, 90% B; 3 min, 75% B; 7 min, 75% B; 8 min, 70% B; 9 min, 70% B; 10 min, 50% B; 12 min, 50% B; 13 min, 25% B; 14 min, 25% B; 16 min, 0% B; 20.5 min, 0% B; 21 min; 90% B; 25 min, 90% B. Autosampler temperature was 4°C, column temperature 30°C and injection volume 3 µL. Samples were injected using electrospray ionization into a QExactive HF orbitrap mass spectrometer (ThermoFisher Scientific) operating in negative ion mode with a resolving power of 75,000 at m/z of 200 and a full scan range of 75–1000. Data were analyzed using the EL-MAVEN software package and specific peaks assigned based on exact mass and comparison with known standards^49^. Extracted peak intensities were corrected for natural isotopic abundance using the R package AccuCor^50^.

### TCA cycle labeling contribution analysis

Total carbon atom fractional contributions were determined by dividing the natural isotope abundance corrected experimental labeling fractions by the number of carbons of each metabolite to determine a single carbon weighted labeling fraction for each metabolite. These were normalized to the serum tracer weighted labeling fraction (presented as 1 in all graphs)^30,51^.

### NAD(H) Quantification by Mass Spectrometry

Each cryo-milled heart sample was extracted on ice with a solution of 3:1 ACN:ddH_2_O, 0.1% NH_4_OH (Millipore Sigma, Burlington, MA, USA) and 0.1 µg/mL carnitine-d9 internal standard (Cambridge Isotope Laboratories, Inc, Tewksbury, MA, USA) pre-chilled at −20 °C to a final tissue concentration of 80 mg/mL. A process blank was created at this time containing only extraction solvent with the internal standard and carried through the extraction process. Samples were vortexed for 30 s, sonicated on ice for 5 minutes, followed by incubation at −20 °C for 15 minutes. Samples were then centrifuged at 20,000 × g for 10 minutes at 4 °C. The supernatant was transferred to PTFE autosampler vials (Agilent Technologies, Inc, Santa Clara, CA, USA) for immediate analysis. A SCIEX 7600 Zeno-ToF with TurboIonSpray ESI source (AB SCIEX LLC, Framingham, MA, USA) coupled to an Agilent 1290 Infinity II HPLC system (Agilent Technologies, Inc, Santa Clara, CA, USA) in positive-ionization mode was used for analysis. Chromatographic separation was achieved using a Waters Atlantis Premier BEH ZHILIC 100 x 2.1 mm column (Waters Corporation, Milford, MA, USA) with Phenomenex Krudkatcher Ultra (Phenomenex, Torrence, CA, USA). Buffers consisted of 99% ACN with 5% ddH2O (buffer B) and 25 mM ammonium carbonate (Sigma-Aldrich, St. Louis, MO, USA) in ddH2O (buffer A). An initial concentration of 99% buffer B was decreased to 85% over 2 minutes, then further decreased to 75% over 3 min, and 60% over 5 min. Next, buffer B was decreased to 40% over 1 minute and held for 1 min. Finally, buffer B was decreased to 1% over 1 min and held for 1 min. Eluents were returned to initial conditions over 0.1 minutes, and the system was allowed to equilibrate for 6.9 minutes between runs. Mass spectrometry analysis was performed by high-resolution multiple reaction monitoring (MRM HR). Source conditions were Curtain gas = 35 psi, CAD gas = 12 psi, Ion source gas 1 = 20 psi, Ion source gas 2 = 30 psi, temperature = 500°C, spray voltage = 5000 V. Metabolites were analyzed with a declustering potential of 50V, and a collision energy of 30V. Data was analyzed in SCIEX Analytics.

### Adult cardiomyocyte isolation

Isolation of adult cardiomyocytes followed previously established protocols^16,39^. Briefly, 12-week-old adult mice were anesthetized with sodium pentobarbital (50 mg/kg) before the heart was excised and attached to an aortic cannula. The heart was then perfused with oxygenated solutions maintained at 37°C and pH 7.3. A 0 mM Ca^2+^ solution was perfused for 5 minutes, followed by a further 15 minutes of perfusion with the same solution containing 1 mg/mL collagenase and 0.1 mg/mL protease. This was succeeded by a 1-minute perfusion with stopping solution (the same solution containing 20% serum and 0.2 mM CaCl_2_), all at a flow rate of 2 ml/min. After removal of the atria, ventricles were teased apart with forceps, gently rocked for 10 minutes, and filtered through a nylon mesh. Following gravity sedimentation, the resulting cells were stored at 37°C in normal HEPES buffered solution before plating as described below. The resulting isolated myocytes displayed a rod-shaped morphology with distinct striations and exhibited no spontaneous contractions.

### ACM culture and physiological media

Human plasma-like media was made in-house as previously published^19^. This enabled the creation of calcium-free media. Glucose or lactate was selectively excluded and replaced with heavy isotope tracers as required. Additionally, the plating media consisted of 100 μg/mL Primocin, 100 units/mL Penicillin-Streptomycin, 10 mM HEPES, 5% dialyzed fetal bovine serum, and 10 mM 2,3-butanedione monoxime. Primary adult cardiomyocytes were isolated (as described above) and enriched through gravity sedimentation in increasing concentrations of Ca^2+^. Following isolation, cells were plated on Laminin-coated Petri dishes or coverslips and allowed to adhere for at least 1 hr in the incubator. Subsequently, cells were transitioned to culture media containing Primocin, Penicillin-Streptomycin, HEPES, Insulin-Transferrin-Selenium, 0.1 mg/mL Bovine serum albumin, and a physiological BSA-conjugated fatty acid mix, as previously described^52^. The cells were cultured in media with 1.2 mM Ca^2+^.

### ACM contractility assay protocol

Contractility assay was performed as previously described^53^. Briefly, the Plexiglas cell bath with a clear glass bottom was mounted on the stage of an inverted microscope (Diaphot, Nikon, Japan), and the ACMs were cultured in a cell super-fusion chamber coated with laminin. Bathing solution in the chamber was maintained at 36 ± 0.3 °C. Cells were field stimulated at a cycle length of 1 s and contractility measured using an inbuilt camera. The bathing solution used contained the following (mM): 126.0 NaCl, 11.0 dextrose, 4.4 KCl, 1.0 MgCl_2_, 1.08 CaCl_2_, and 24.0 HEPES titrated to pH 7.4 with 1 M NaOH. The pipette solution used for recording APs contained the following (mM): 110.0 KCl, 5.0 NaCl, 5.0 MgATP, 5.0 phosphocreatine, 1.0 NaGTP, 10.0 HEPES titrated to pH 7.2 with 1 M KOH.

### Mitochondrial isolation

One whole mouse heart was minced in ice-cold mitochondrial isolation medium (MIM) buffer [300 mM sucrose, 10 mM Hepes, 1 mM EGTA, and bovine serum albumin (BSA; 1 mg/mL) (pH 7.4)] and gently homogenized with a Teflon pestle. Samples were centrifuged at 800 *x g* for 10 min at 4°C. The supernatants were then transferred to fresh tubes and centrifuged again at 1,300 x g for 10 min at 4°C. To achieve the mitochondrial fraction (pellet), the supernatants were again transferred to new tubes and centrifuged at 9,000 x g for 10 min at 4°C. The final mitochondrial pellets were resuspended in MIM buffer for experimental use. This crude mitochondrion preparation was used for the metabolomics and respirometry experiments. For the immunoblotting, immunoprecipitation, and proteomics experiments, these preparations were further purified by ultracentrifugation on a two-step (1.33 M, 1.55 M) sucrose gradient. Gradients were centrifuged at 4°C for 1h at 22,500 rpm. Mitochondria formed a compact, brown-colored band at the interface. An 18-gauge needle was used to carefully recover the mitochondrial band for further analysis.

### Mitochondrial respiration measurements

Mitochondrial O_2_ utilization was measured using the Oroboros Oxygraph-2K system (Innsbruck, Austria). Following BCA assay, freshly isolated mitochondria (25 µg) were added to the respirometry chambers containing 2mL of assay buffer Z (MES potassium salt 105 mM, KCl 30 mM, KH_2_PO_4_ 10 mM, MgCl_2_ 5 mM, BSA 1 mg/ml). Respiration on lactate or pyruvate was measured with the following additional substrates: 0.5 mM pyruvate or 0.5 mM lactate with 1 mM ADP and 500 µM malate. Inhibitors of pyruvate and lactate metabolism were added to the buffer in the following concentrations: CPI-613 (PDH and αKGDH inhibitor) 240 μM, GSK 2837808A (pan-LDH inhibitor) 8 μM, and 7ACC2 (MCT1 inhibitor) 10 μM or DMSO control.

### In vitro metabolite tracing

ACMs were plated as described in HPLM-FA media. For tracing experiments, glucose- or lactate-free HPLM-FA media was made in-house, and [U-^13^C]glucose or [U-^13^C]lactate was added to the media. Cells were incubated for 4 hrs. Isolated mitochondria were treated identically as reactions were carried out at 37°C in HPLM-FA media with appropriate ^13^C-tracers and quenched with −80°C cold methanol. Cell plates were washed with ice cold PBS 2x and then 80:20 methanol:water was added at 60x of the PCV (packed cell volume) (MidSci, TP87005) count. The resulting mixture was incubated on dry ice, scraped, collected into a microfuge tube, vortexed, rested on dry ice for 5 minutes and centrifuged at 16000 × g for 10 minutes. Supernatant was placed into a fresh tube which was then centrifuged again at 16000 x for 10 minutes. The supernatant was placed in an MS tube (Agilent 5188-2788) for downstream analysis.

### MCT1 imaging studies

Isolated adult cardiomyocytes were transferred to a 1.5 mL microcentrifuge tube and fixed for 15 min with 4% paraformaldehyde in 1× PBS followed by permeabilizing for 7 minutes with 0.1% Triton X-100 in 1× PBS. Fixed samples were washed three times with 1× PBS and blocked overnight with 5% bovine serum albumin (BSA) in 1× PBS at 4°C. Immunostaining was performed with MitoTracker Red CMXRos (catalog no.: M7512; Invitrogen) at room temperature for 15 min. prior to fixing, and MCT1 (catalog no.: 365501; SCBT), SLC25A6 (catalog no.: 154007; Abcam) or TOMM20 primary antibodies (catalog no.: 42406; CST) at room temperature for 1 hr. Samples were then washed four times with 1× PBS and incubated at room temperature for 1 hr with Alexa Fluor 568 goat anti-rabbit secondary antibody (catalog no.: A11011; Life Technologies) or Alexa Fluor 488 goat anti-mouse secondary antibody (catalog no.: A11017; Life Technologies). After incubation, the samples were washed four times with 1× PBS and mounted in Vectashield Vibrance antifade mounting media with 40,6-diamidino-2-phenylindole (DAPI; catalog no.: H-1800; Vector Laboratories). Images were acquired with a Zeiss LSM 880 confocal microscope with Airyscan. Intensity profiles and mean intensity values were measured at each pixel along the line by manually drawing a 1-pixel width line along the long axis of a mitochondria using ImageJ (National Institutes of Health). Scale bars of 10 μm were added to merged images using ImageJ.

### Immunoblotting

Samples were washed with PBS and lysed in RIPA buffer (50 mM Tris, 150 mM NaCl, 0.1% SDS, 0.5% sodium deoxycholate, 1% NP-40) containing protease and phosphatase inhibitors. Protein concentrations were determined using the Pierce BCA Protein Assay Kit. Samples were combined with 4x sample loading buffer and heated at 95°C for 5 minutes. A total of 20 μg of protein lysate was separated on an SDS polyacrylamide gel following standard procedures at 20 mA per gel, then transferred to a 0.45 μm nitrocellulose membrane (GE Healthcare) using a Mini Trans-blot module (Bio-Rad) at a constant voltage of 100 V for 2 hrs. The membrane was blocked with 5% non-fat milk (Serva) in Tris-buffered saline with 0.05% Tween 20 (TBS-T) overnight at 4°C and then incubated for 3 hrs to overnight in 5% non-fat milk or 5% bovine serum albumin (Sigma) in TBS-T with primary antibodies against MPC1 (Cell Signaling, 1:1000), MPC2 (Cell Signaling, 1:1000), VDAC (Cell Signaling, 1:5000), MCT1 (Santa Cruz, 1:500, Proteintech, 1:1000), LDHA (Proteintech, 1:1000), LDHB (Proteintech, 1:1000), COX IV (Cell Signaling, 1:1000), ATP5A (Cell Signaling, 1:1000), or TOM20 (Cell Signaling, 1:1000). The membrane was then washed with TBS-T and incubated with the appropriate fluorophore-conjugated secondary antibody (Rockland Immunochemical, 1:10000) in 1% non-fat milk/TBS-T for 30 minutes. After a final wash with TBS-T, fluorescence was detected using the Odyssey CLx imaging system (LI-COR Biosciences).

### MCT1 Immunoprecipitation (IP)

Mitochondrial preparations were washed twice with ice-cold PBS before being pelleted by centrifugation at 1200 rpm for 3 minutes. The pellets were lysed in 50µL of RIPA lysis buffer on ice for 10–20 minutes. Lysates were then centrifuged at 16,100 × g for 10 minutes, and the supernatants were transferred to fresh tubes. For each condition, immunoprecipitations (IP) were performed as follows: Experimental IP: Anti-MCT1 mouse antibody was added to the supernatant at a ratio of 1.5–2 µL antibody per sample. Control IP: Normal mouse IgG (Rb IgG) was diluted to match the concentration of the experimental antibody and added to a separate sample of lysate. Both IP reactions were incubated overnight at 4°C with rotation. The next day, samples were washed 3 times with lysis buffer. For each IP, 25–35 µL of pre-washed beads were added to the samples, and incubated with rotation for 2 hrs at 4°C. Beads were then collected by a magnet, and washed three times with lysis buffer. After washing, beads were eluted and immunoprecipitated proteins separated by SDS-polyacrylamide gel electrophoresis and either immunoblotted for MCT1 or band at correct size was excised and directly sent for proteomic analysis. Gel bands were excised, and their protein content was digested in-gel with trypsin^54,55^. The extracted peptides were desalted via StageTip, dried using vacuum centrifugation, and reconstituted in 5% acetonitrile with 5% formic acid for LC-MS/MS processing^56^.

### Proteomics (Tandem Mass Spectrometry)

Mass spectrometric data were collected on Orbitrap Fusion Lumos instruments coupled to a Proxeon NanoLC-1200 UHPLC. The 100 µm capillary column was packed with 35 cm of Accucore 150 resin (2.6 μm, 150Å; ThermoFisher Scientific) at a flow rate of 360 nL/min. The scan sequence began with an MS1 spectrum (Orbitrap analysis, resolution 60,000, 350-1350 Th, automatic gain control (AGC) target 100%, maximum injection time of 118ms). Data were acquired for 60 minutes per sample. The hrMS2 stage consisted of fragmentation by higher energy collisional dissociation (HCD, normalized collision energy 36%) and analysis using the Orbitrap (AGC 200%, maximum injection time 60 ms, isolation window 1.2 Th, resolution 7.5K). Data were acquired using the FAIMSpro interface the dispersion voltage (DV) set to 5,000V, the compensation voltages (CVs) were set at −40V, −60V, and −80V, and the TopSpeed parameter was set at 1 sec per CV. Mass spectra were processed using a Comet-based in-house software pipeline. MS spectra were converted to mzXML using a modified version of ReAdW.exe. Database searching included all entries from the human UniProt database (downloaded November 2021), which was concatenated with a reverse database composed of all protein sequences in reversed order. The digest was set to semi-tryptic. Searches were performed using a 50 ppm precursor ion tolerance and the product ion tolerance was set to 0.02 Th. Oxidation of methionine residues (+15.9949 Da) was set as a variable modification. PSM filtering was performed using a linear discriminant analysis, as described previously^57^, while considering the following parameters: XCorr, ΔCn, missed cleavages, peptide length, charge state, and precursor mass accuracy. Peptide-spectral matches were identified, quantified, and collapsed to a 1% FDR and then further collapsed to a final protein-level false discovery rate (FDR) of 1%.

### Proteinase K protection assay

Briefly, 50 μg of purified mitochondria were added to various assay buffers: SEM (250 mM Sucrose, 10 mM MOPS/KOH pH 7.4, 1 mM EDTA), EM (10mM MOPS/KOH pH 7.4, 1 mM EDTA), or SEM + 1% TX-100 (SEM supplemented with 1% Triton X-100). After brief vortexing, the samples were then incubated on ice for 5 minutes. Next, Proteinase K (PK) was added to a final concentration of 25 μg/mL to half of the samples, then vortexed and incubated on ice for 10 minutes. Phenylmethylsulfonyl fluoride (PMSF) was added to stop PK digestion (final concentration 2 mM). Samples were vortexed again and incubated for another 10 minutes on ice. SEM and EM samples were centrifuged at 20,000 x g for 10 minutes at 4°C. TX-100 samples were precipitated with trichloroacetic acid (TCA) (final concentration 20%). 15 μL of each sample was loaded onto SDS-PAGE for analysis.

### Quantitative PCR Analysis

Total RNA from mouse hearts was isolated using RNeasy Mini Kits (Qiagen), according to the manufacturer’s instructions. Next, cDNA was synthesized using a cDNA Reverse Transcriptase Kit (New England Biolabs). TaqMan-based real time quantitative polymerase chain reactions (qRT-PCR) were then performed using a QuantStudio 7 Pro Real-Time PCR System (ThermoFisher). The housekeeping gene *Vinculin* was used as an internal control for cDNA quantification and normalization of gene amplified products.

### RNA sequencing and data analysis

RNA was isolated from murine ventricular tissue samples using the miRNeasyMini Kit (QIAGEN). Total RNA samples (100-200 ng) were hybridized with Ribo-Zero Gold (Illumina) to substantially deplete cytoplasmic and mitochondrial rRNA from the samples. Stranded RNA sequencing libraries were prepared as described using the Illumina TruSeq Stranded Total RNA Library Prep Gold kit (20020598) with TruSeq RNA UD Indexes (20022371). Purified libraries were qualified on an Agilent Technologies 2200 TapeStation using a D1000 ScreenTape assay (cat# 5067-5582 and 5067-5583). The molarity of adaptor-modified molecules was defined by quantitative PCR using the Kapa Biosystems Kapa Library Quant Kit (cat#KK4824). Individual libraries were normalized to 1.30 nM, were chemically denatured and applied to an Illumina NovaSeq flow cell using the NovaSeqXP chemistry workflow (20021664). Following transfer of the flowcell to an Illumina NovaSeq instrument, a 2 x 51 cycle paired end sequence run was performed using a NovaSeqS1 reagent Kit (20027465).

RNA-seq analysis was conducted with the High-Throughput Genomics and Bioinformatic Analysis Shared Resource at Huntsman Cancer Institute at the University of Utah. The mouse GRCm38 FASTA and GTF files were downloaded from Ensembl release 96 and the reference database was created using STAR version 2.7.0f with splice junctions optimized for 50 base-pair reads (Dobin et al., 2013). Optical duplicates were removed from the paired end FASTQ files using BBMap’s Clumpify utility (v38.34) (https://sourceforge.net/projects/bbmap) and reads were trimmed of adapters using cutadapt 1.16 (Martin, 2011). The trimmed reads were aligned to the reference database using STAR in two-pass mode to output a BAM file sorted by coordinates. Mapped reads were assigned to annotated genes in the GTF file using featureCounts version 1.6.3 (Liao et al., 2014). The output files from cutadapt, FastQC, Picard CollectRnaSeqMetrics, STAR, and featureCounts were summarized using MultiQC to check for sample outliers (Ewels et al., 2016). Differentially expressed genes with at least 95 read counts across all samples were identified using DESeq2 version 1.24.0 (Love et al., 2014). Differentially expressed genes were then identified with a *q*-value > 0.05. Enriched diseases and/or biological functions analysis were performed by Ingenuity Pathway Analysis (IPA) software (QIAGEN Bioinformatics, Redwood City, CA). Differentially expressed genes from [4 comparison groups] were uploaded into IPA. Analysis ready molecules were confined to mouse and heart genes with a q-value > 0.05 and log2FoldChange > |0.074|, equivalent to a fold change of 5%. Fischer’s exact test was used to calculate a *p*-value determining the probability that each biological function and/or disease assigned to these data sets were due to chance alone. Significance threshold was set at *p-*value <0.05 and z > |2|.

### In vitro hypoxia/reoxygenation injury

Imaging was performed as described previously^58^. Briefly, isolated cardiomyocytes were suspended in HPLM-FA media and placed in an airtight chamber at 37°C. To stimulate ischemic conditions, the chamber was flushed with a gas mixture containing 95% N_2_ and 5% CO_2_ to deplete the oxygen in the chamber. The chamber was then sealed, and the cells left for 2 hrs at 37 °C. After 2 hrs, the cells were removed from the chamber and placed under normal cell culture conditions (37°C, room air supplemented with 5% CO_2_) for an additional 2 hrs to reoxygenate and model reperfusion. Control cells were suspended in HPLM media and kept at 37°C, in room air with 5% CO_2_ for 4 hrs. For imaging, the cells were loaded with either 20 nM of TMRM (Thermo Fisher), 5 μM MitoSox (Thermo Fisher), or 5 μM X-Rhod1 (Thermo Fisher) to measure mitochondrial membrane potential, mitochondrial ROS levels and mitochondrial Ca^2+^ levels, respectively. The cells were then placed onto glass-bottomed dishes (MatTek, MA) and imaged using a Leica SP8 confocal microscope (Deer Park, IL). All three dyes were measured with an excitation and emission at 548nm/ 574 nm. All cells were imaged at the same power and gain settings and images were collected within 30 minutes after removal from incubator following simulated reperfusion. Image analysis was performed using ImageJ^59^.

### Seahorse assay

The Seahorse Mitochondrial Stress Test was performed following the manufacturer’s instructions on a Seahorse XF Pro Analyzer. Adult cardiomyocytes were plated in a 96-well seahorse plate in HPLM media without BSA-conjugated fatty acids. The Seahorse assay was performed using 3 μM Oligomycin, 1 μM FCCP and 1 μM Rotenone/1 μM Antimycin A, with a standard protocol of three measurement cycles for each phase (2 minutes mixing, 3 minutes waiting and 3 minutes measuring). After the assay, data were analyzed in the Seahorse WAVE software through the XF Mito Stress Test Report.

### Statistical Analysis

We used GraphPad Prism software (v10.3.1) for all statistical analysis. To determine the if our data was normally distributed, we used the Shapiro-Wilk and D’Agostino-Pearson omnibus tests. For in vivo experiments, we utilized Grubb’s method (Extreme Studentized Deviate; ESD: to detect one outlier) and ROUT (Robust Outlier Detection: to detect multiple outliers) testing was to expose any significant outliers (p<0.05) in our data. If outliers were detected they were removed subsequent statistical analysis. For all of our data we present it as the mean ± standard error of the mean (SEM) and applied 2-tailed tests for all comparisons. When analyzing two discrete groups (such as *Mct1^fl/fl^* vs. *Mct1^iCKO^*), we employed an unpaired Student’s *t*-test for normally distributed data, if the data was non-parametric, we used the Mann-Whitney U test. Multiple comparisons correction for unpaired t-tests was performed using the Holm-Sídák method for adjusted p-values. For datasets with more than two variables one and two-way ANOVA was used with Tukey’s multiple comparison test. For evaluating serial echocardiographic data and multiple group comparisons, we used a repeated measures ANOVA (mixed-effects model) with a Geisser-Greenhouse correction, followed by a Tukey’s multiple comparison test for individual comparisons. We set an α level of p<0.05 *a priori* and any value below this setpoint was considered statistically significant.

## Data availability

All data presented in our manuscript is available from the corresponding authors upon reasonable request. All materials used in this study, including mouse models and other resources, can be shared upon request of the corresponding authors. Distribution of these materials may require completion of a materials transfer agreement (MTA) to ensure intellectual property protections. We are committed to fostering collaboration and knowledge sharing within the scientific community to ensure adherence to ethical and institutional guidelines.

## Acknowledgements

We thank the Nora Eccles Harrison Cardiovascular Research and Training Institute and the Nora Eccles Harrison Treadwell Foundation for their support of this research project. We also thank the H.A. Edna Benning Society at the University of Utah for funding provided to S.G.D. We thank the University of Utah Department of Biochemistry and the Diabetes and Metabolism Research Center for support to G.S.D. We acknowledge funding from the following sources: the National Institutes of Health (NIH) under Ruth L. Kirschstein National Research Service Award T32HL007576 from the National Heart, Lung and Blood Institute to J.R.V, 5T32DK091317 from the National Institute of Diabetes and Digestive and Kidney Diseases to J.N.V., awards R01HL135121 (NIH), 1R01HL166513 (NIH), I01 CX002291 (U.S. Department of Veterans Affairs), I01BX006306-01 (U.S. Department of Veterans Affairs) and 16SFRN29020000 (American Heart Association) to SGD, award K99HL168312 to A.A.C., award R01HL141353 and R01HL165797 to D.C., award 3R35GM131854-04 to J.R. from the National Institute of General Medicine Sciences, award 834544 to D.R.E and award 1019351 to T.S.S from the American Heart Association. We would also would like to thank the Burroughs Wellcome Fund for Postdoctoral Diversity Enrichment Program award to L.C.R. Research reported in this publication utilized the High-Throughput Genomics and Cancer Bioinformatics Shared Resources at the University of Utah Huntsman Cancer Institute, which is supported by the National Cancer Institute of the National Institutes of Health under Award Number P30CA042014. Lastly, J.R. is an investigator of the Howard Hughes Medical Institute. We thank the University of Utah Metabolic Phenotyping Core and the members of the Drakos, Rutter, and Ducker labs for their assistance and helpful discussions. The content of this manuscript is solely the responsibility of the authors and does not necessarily represent the official views of the NIH.

## Author information

These authors contributed equally: Ahmad A. Cluntun, Jesse N. Velasco-Silva, and Joseph R. Visker. The order of the co-first authors was assigned by mutual agreement.

## Ethics declarations

Animal experiments were conducted in accordance with the institutional guidelines for the care and use of laboratory animals. Our protocols were reviewed and approved by the Institutional Animal Care and Use Committee (IACUC) are the University of Utah (21-02003). For experiments involving the use of human myocardial tissue, our ethical approval was obtained from the Institutional Review Board (IRB) at the University of Utah (00030622) and all patients provided written informed consent prior to their inclusion in the study.

## Supplementary information

Supplemental Data File A: Video of cardiac contractility of ACMs cultured in HPLM-FA

Supplemental Data File B: Metabolomics data from Ang/PE treated *Mct1^iCKO^* hearts

Supplemental Data File C: RNAseq data from Ang/PE treated *Mct1^iCKO^* hearts

Supplemental Data File D: Qiagen IPA analysis of differentially expressed pathways from Ang/PE treated *Mct1^iCKO^* hearts

Supplemental Data File E: Source data used to generate figure panels from Fig.1-4 and Extended Data 1-11.

## Source data

Our transcriptomics data has been deposited and are publicly available on the NCBI-NIH Sequence Read Archive (SRA) and Gene Expression Omnibus (GEO) repository under the accession number of GSE276036. Source data for all panels is included as Supplemental Date File E.

## Author Contributions

This work was conceived of by A.A.C., S.G.D., J.R. and G.S.D. and they wrote the paper with input from all of the other authors. In vivo stable isotope tracing experiments were carried out by J.N.V. with assistance from J.E.K., H.K.L., and C.E.S. In vitro cellular, biochemical, and imaging assays were carried out by A.A.C. with help from M.J.L., K.F., L.C-R., A.G.T., C.N.C., J.C., M.Y.J., A.J.B, A.J.N-P., J.T.M., T.Y., D.R.E., S.A.B., D.K.A.R. and D.C. Functional cardiac studies in mice were led by J.R.V. with assistance from T.S.S., R.H. Additional mass spectrometry experiments were performed and analyzed by A.A.C., Q.P., J.L.C., C.E.W., S.P.G., J.A.P., J.E.C. and G.S.D. Animal model generation was led by J.L., S.N.. J.D.R. and W.I.S. Q.L. performed transcriptomics analysis. D.M.M. provided insightful input on experimental design and interpretation. A.A.C., J.R.V. and J.N.V analyzed data with assistance from G.S.D. S.G.D., J.R., and G.S.D. managed the funding, resources, and reagents necessary for this project.

## Extended Data Figure Legends

**Extended Data Figure 1.**
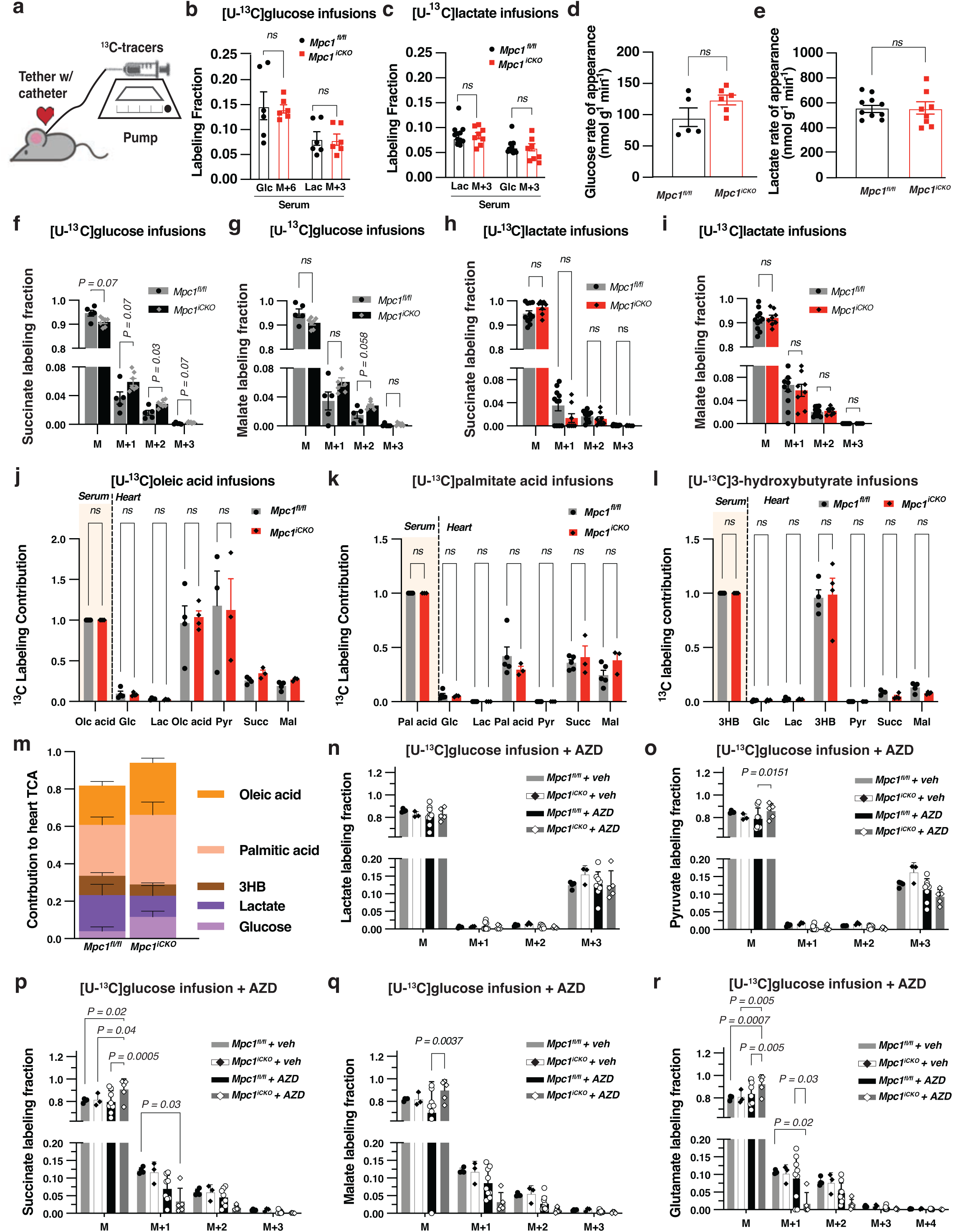
Cardiac fuel usage in *Mpc1^iCKO^* mice. **a,** Schematic showing stable awake infusion experiment setup for mouse. **b,** Serum [U-^13^C]glucose labelling fraction after 3 hrs infusion in fasted mice (*Mpc1^fl/fl^ n=6*, *Mpc1^iCKO^ n=6*). **c**, Serum [U-^13^C]lactate labelling fraction after 2 hours infusion in fasted mice (*Mpc1^fl/fl^ n=12*, *Mpc1^iCKO^ n=8*). **d,** Glucose and **e,** lactate rate of appearance calculated from labelling fractions in **a**, **b**. **f,** Succinate and **g,** malate heart ^13^C labelling fractions from [U-^13^C]glucose infusions (*Mpc1^fl/fl^ n=6*, *Mpc1^iCKO^ n=6*) **h,** Succinate and **i,** malate heart ^13^C labelling fractions from [U-^13^C]lactate infusions (*Mpc1^fl/fl^ n=12*, *Mpc1^iCKO^ n=8*). **j,** Serum normalized ^13^C labeling of heart lactate, pyruvate and TCA cycle metabolites from [U-^13^C]oleic acid infusions in *Mpc1^iCKO^* (*n=3*) and *Mpc1^fl/fl^* (*n=4*) mice. **k,** Serum normalized ^13^C labeling of heart lactate, pyruvate and TCA cycle metabolites from [U-^13^C]palmitic acid infusions in *Mpc1^iCKO^* (*n=3*) and *Mpc1^fl/fl^* (*n=5*) mice. **l,** Serum normalized ^13^C labeling of heart lactate, pyruvate and TCA cycle metabolites from [U-^13^C]3-hydroxybutyrate infusions in *Mpc1^iCKO^*(*n=4*) and *Mpc1^fl/fl^* (*n=4*) mice. **m** Calculated direct circulating nutrient contribution to cardiac TCA cycle metabolism (based on succinate and malate labelling data) in *Mpc1^fl/fl^* and *Mpc1^iCKO^* animals. **n-r,** Isotope corrected ^13^C labeling fractions of glycolytic and TCA cycle metabolites from *Mpc1^fl/fl^* and *Mpc1^iCKO^* mice treated with either AZD3965 (AZD) or vehicle (Veh) and infused with [U- ^13^C]glucose (*Mpc1^fl/fl^* Veh *n=4*, *Mpc1^iCKO^* Veh *n=3*, *Mpc1^fl/fl^* AZD *n=8*, *Mpc1^iCKO^ n*=5). Glc: glucose; Lac: lactate; Succ: succinate; Pyr: pyruvate; Mal: Malate. All data represent mean ± SEM. ^∗^p < 0.05, ^∗∗^p < 0.01, ^∗∗∗^p < 0.001, ^∗∗∗∗^p < 0.0001, determined by unpaired t tests (b-e), multiple comparisons corrected unpaired t-tests (j-l), or two-way ANOVA with multiple comparisons correction (Tukeys) (n-r). ns, not significant (p>0.05).

**Extended Data Figure 2.**
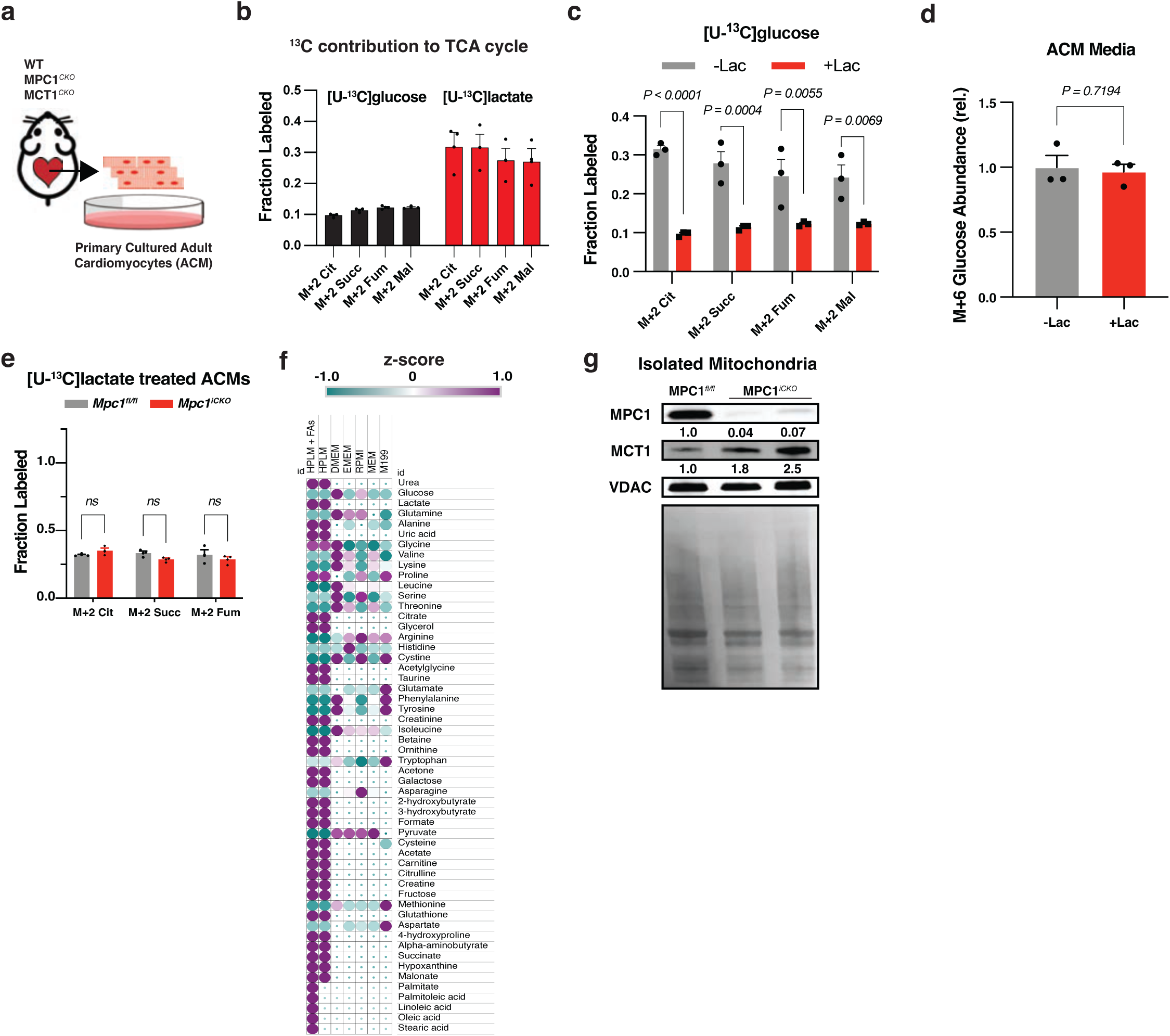
Adult cardiomyocytes preferentially metabolize lactate over glucose. **a**, Schematic of ACM isolations from cardiac specific knockout mice. **b**, ^13^C-enrichment of TCA cycle intermediates from ACMs treated with [U-^13^C]glucose (black) or [U-^13^C]lactate (red), *n=3*. **c**, ^13^C-enrichment of TCA cycle intermediates from ACMs treated with [U-^13^C]glucose in in the presence (gray) or absence (black) of lactate*, n=3*. **d**, Relative abundance of M+6 Glucose in the media after 4 hrs in the presence or absence of lactate, *n=3*. **e**, ^13^C-enrichment of TCA cycle intermediates in ACMs harvested from *MPC1^fl/fl^* or *MPC1^iCKO^* mice cultured with [U-^13^C]lactate, *n=3*. **f**, Heatmap of metabolites present in the formulation of HPLM-FA compared to other standard cell culture medias (clustered by z-scores from log_2_ concentrations). **g**, Representative immunoblot of MPC1, MCT1 and VDAC in mitochondria isolated from *Mpc1^fl/fl^* or *Mpc1^iCKO^* cardiac tissue and ponceau stain of the blot, 4 weeks post induction. All data represent mean ± SEM. Significance determined by multiple comparisons corrected unpaired two-tailed t-tests, or two-tailed t-tests. ns, not significant (p>0.05).

**Extended Data Figure 3.**
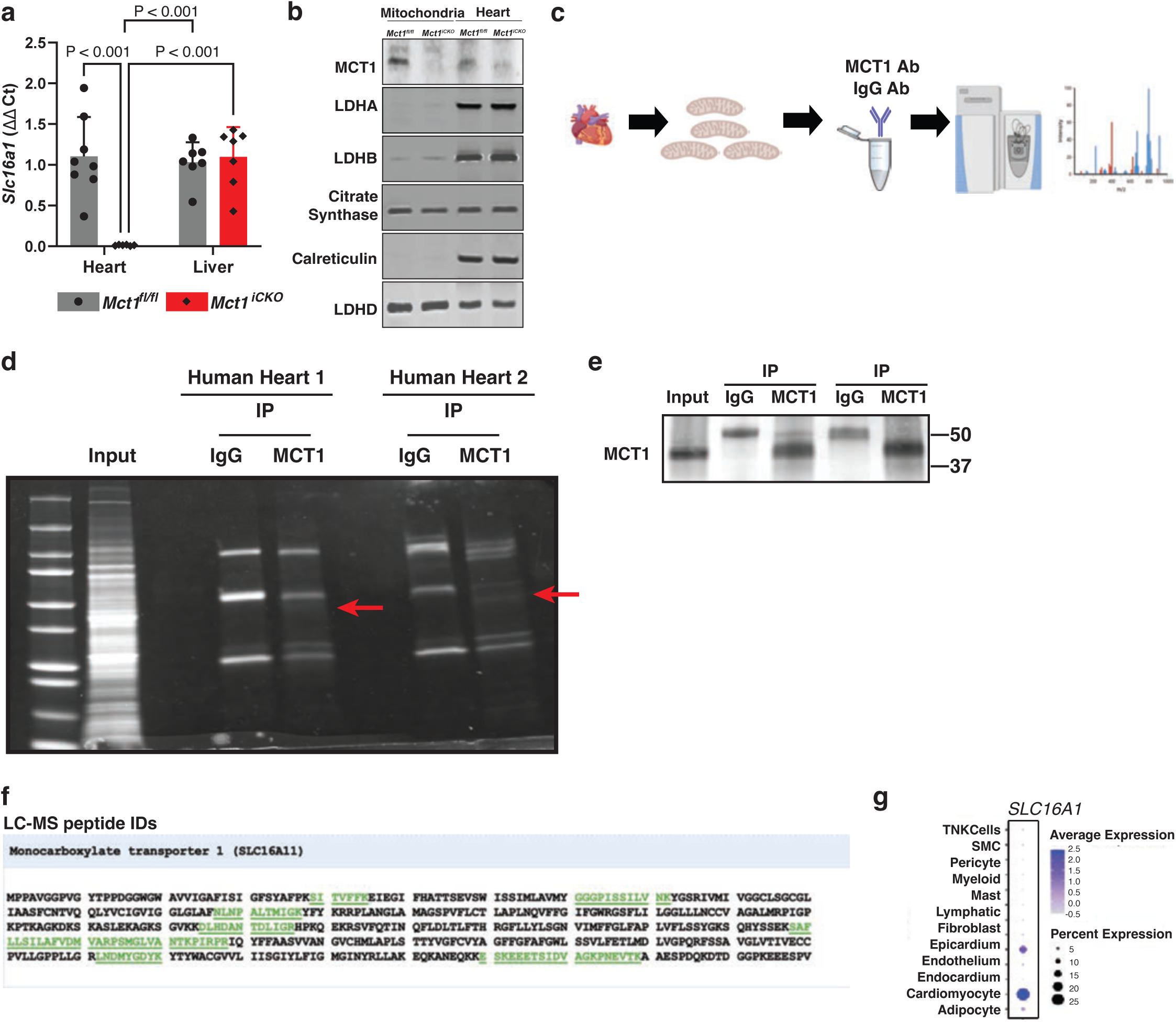
Identification of MCT1 from purified human cardiac mitochondria. **a,** qPCR measurement of *Slc16a1* (Mct1) gene expression in heart and liver lysates from control (*Mct1^fl/fl^*) and cardiac-specific knockout (*Mct1^iCKO^*) mice 4 weeks post-induction (*Mct1^fl/fl^ n=8, Mct1^iCKO^ n=7*). All data represent mean ± SEM. Significance determined by two-way ANOVA with Tukeys multiple comparison. **b,** Representative immunoblot analysis of MCT1, LDHA, LDHB, citrate synthase, calreticulin, and LDHD protein in heart lysates and purified mitochondria from *Mct1^fl/fl^* and *Mct1^iCKO^* mice 4 weeks post-induction. **c,** Schematic of the experimental design. Mitochondria were purified from human patient cardiac tissue, proteins were extracted from these mitochondria and immunoprecipitated by either an MCT1 antibody or an anti-IgG antibody. Eluted immunoprecipitants were run on a SDS Page gel and bands at the correct size were cut out and sent for proteomic analysis. **d**, Representative image of Coomassie blue stained gel of two unique human hearts, and **e**, corresponding immunoblot of MCT1 of the IP experiment. **f**, Sequence coverage (peptides) of human MCT1 from LC-MS/MS measurements of protein lysates from IP pulldown in b. **g,** Expression of SLC16A1 transcripts from single-nuclei data from donor hearts mapped to cell types. Data replotted from reference 25.

**Extended Data Figure 4.**
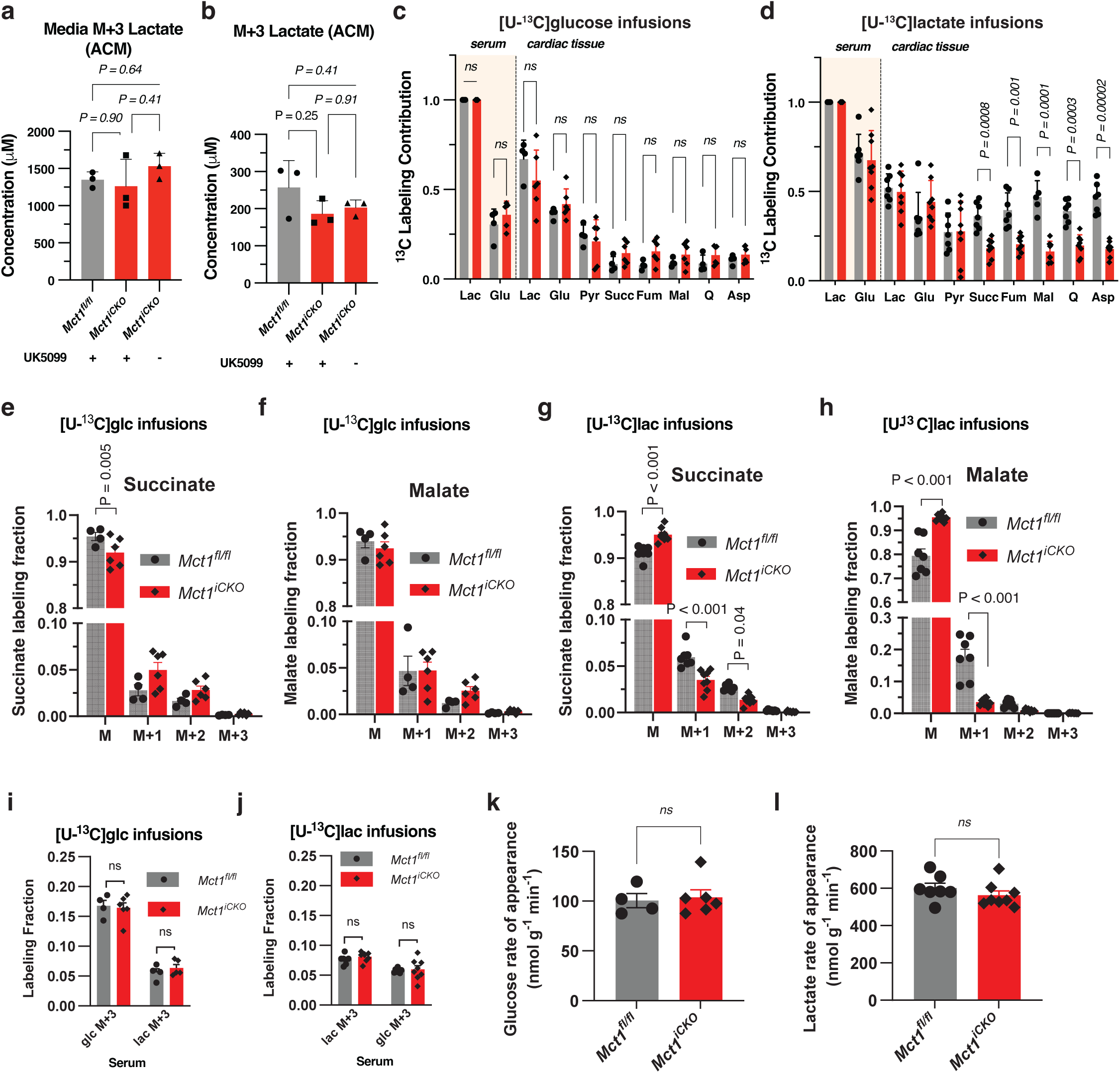
Cardiac fuel usage in *Mct1^iCKO^* mice. **a**, Media M+3 lactate concentration upon UK5099 treatment from ACMs isolated from *Mct1^fl/fl^* and *Mct1^iCKO^*hearts and cultured in [U-^13^C]lactate (*n=3*). **b,** Intracellular M+3 lactate concentration upon UK5099 treatment in cultured ACMs from a (*n=3*). **c,** Heart ^13^C labeling of glucose, lactate, pyruvate and TCA cycle metabolites from [U-^13^C]glucose infusions in *Mct1^fl/fl^* (*n=4*), or *Mct1^iCKO^* (*n=6*) mice. **d**, Heart ^13^C labeling of glucose, pyruvate, lactate and TCA cycle metabolites from [U-^13^C]lactate infusions in *Mct1^fl/fl^* (*n=7*) or *Mct1^iCKO^* (*n=8*) mice. **e,** Succinate and **f,** malate heart ^13^C labelling fractions from [U-^13^C]glucose infusions (*Mct1^fl/fl^ n=4*, *Mct1^iCKO^ n=6*). **g,** Succinate and **h,** malate heart ^13^C labelling fractions from [U-^13^C]lactate infusions (*Mct1^fl/fl^ n=7*, *Mct1^iCKO^ n=8*). **i,** Serum [U- ^13^C]glucose labelling fraction after 3 hours infusion in fasted mice (*Mct1^fl/fl^ n=4*, *Mct1^iCKO^ n=6*). **j**, Serum [U-^13^C]lactate labelling fraction after 2 hours infusion in fasted mice (*Mct1^fl/fl^ n=7*, *Mct1^iCKO^ n=8*). **k,** Glucose and **l,** lactate rate of appearance calculated from steady state labelling fractions in f, g. Glc: glucose; Lac: lactate; Pyr: pyruvate; Succ: succinate; Fum: fumarate; Mal: Malate; Q: glutamate; Asp: aspartate. All data represent mean ± SEM. Significance determined by unpaired t-tests (i,j), one-way ANOVA with Dunnett’s multiple comparison test (a,b), multiple comparisons corrected unpaired two-tailed t-tests (c-h), two-way ANOVA with Tukeys multiple comparison (b), ns, not significant (p>0.05).

**Extended Data Figure 5.**
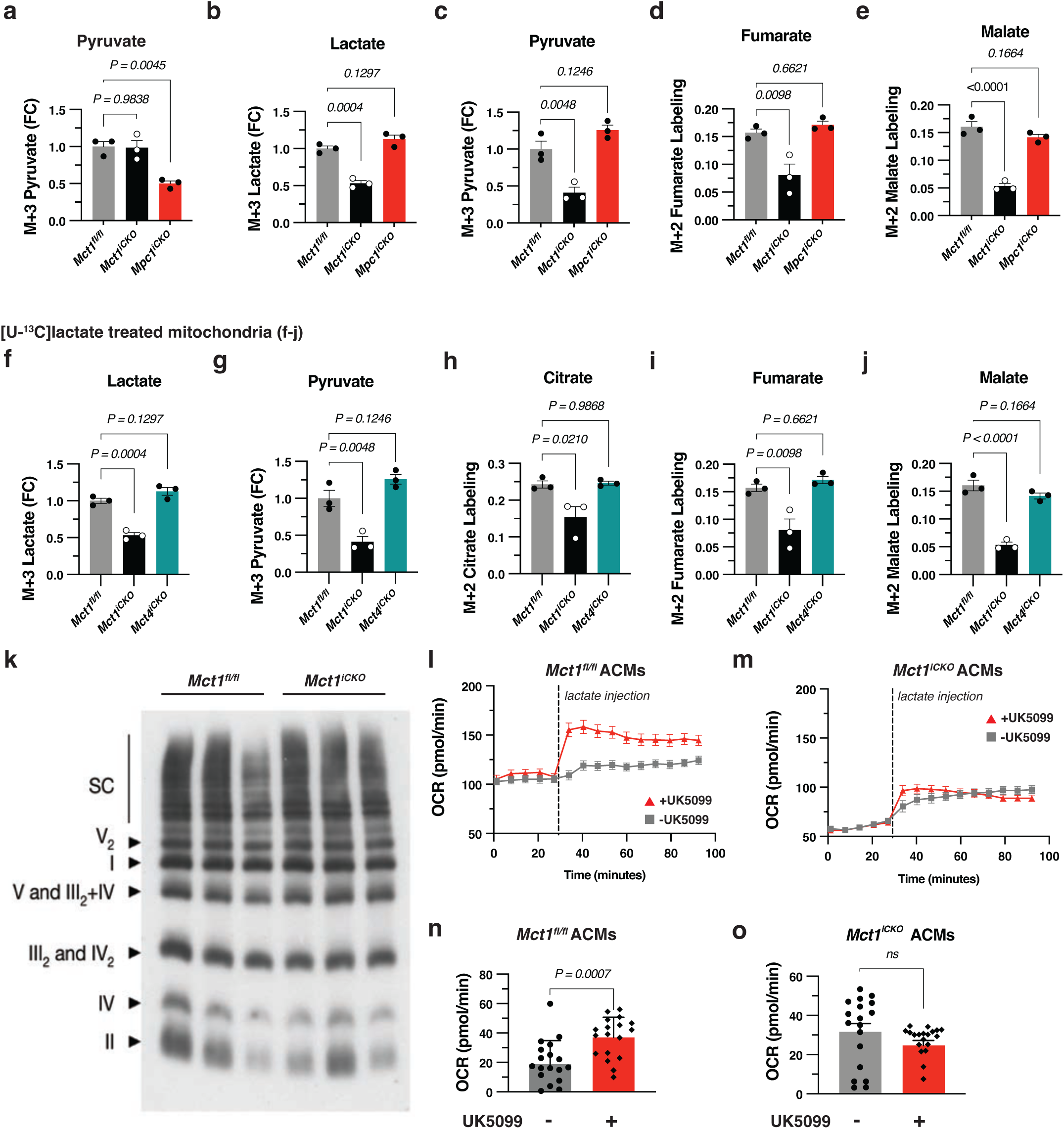
Mitochondrial metabolism and respiration on lactate depends on MCT1. **a**, Relative abundance of M+3 pyruvate in mitochondria from the indicated genotypes treated with [U-^13^C]pyruvate. **b**, Relative abundance (fold change, FC) of M+3 lactate in mitochondria from the indicated genotypes treated with [U-^13^C]lactate, and labeling fraction of downstream metabolites, M+3 pyruvate (**c**), M+2 fumarate (**d**), and M+2 malate (**e**), *n=3*. **f**, Relative abundance (FC) of M+3 lactate in mitochondria from indicated genotypes, treated with [U-^13^C]lactate, and downstream metabolites, M+3 pyruvate (**g**), M+2 citrate (**h**), M+2 fumarate (**i**), and M+2 malate (**j**), *n=3*. **k**, Native blue stain gel of mitochondria isolated from *Mct1^fl/fl^*, and *Mct1^iCKO^* hearts, *n=3*. **l**, Seahorse oxygen consumption assay of ACMs from lactate pre-treated with UK5099 or Vehicle in *Mct1^fl/fl^* or, **m**, *Mct1^iCKO^* ACMs. **n**, Oxygen consumption rates of *Mct1^fl/fl^* ACMs pretreated with UK5099 or Vehicle, and, **o**, in *Mct1^iCKO^* ACMs. All data represent mean ± SEM. Significance determined by one-way ANOVA with Dunnett’s multiple comparison test and unpaired two-tailed t-tests (n,o). ns, not significant (p>0.05).

**Extended Data Figure 6.**
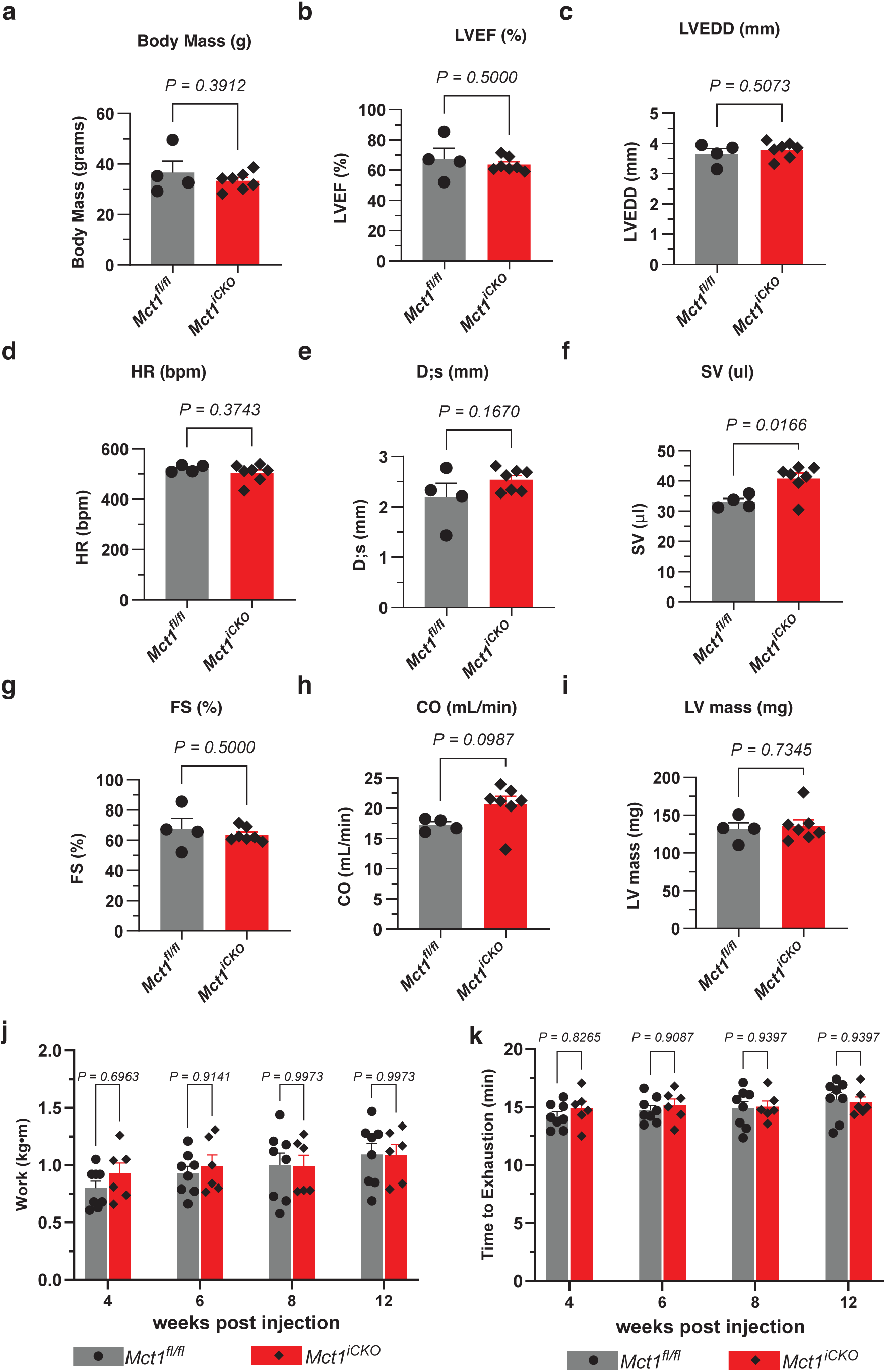
Normal cardiac function and exercise tolerance in 1-year old *Mct1^iCKO^*mice. Echocardiography was performed on mice (*Mct1^fl/fl^ n=4*, and *Mct1^iCKO^ n=7*) older than 1 year of age. **a,** Body mass (g: grams). **b**, Left ventricular ejection fraction (LVEF: %). **c**, Left ventricular end diastolic diameter (LVEDD: mm). **d**, Heart rate (HR: bpm). **e**, Left ventricular end systolic diameter (D;s: mm). **f**, Stroke volume (SV: μL). **g**, Fractional shortening (FS: %). **h**, Cardiac output (CO: mL/min). **i**, absolute left ventricular mass (LV mass: mg). **j** and **k**, Mice were subjected to exercise tolerance tests on a small rodent treadmill and work (kg/m) and time to exhaustion (min) was calculated at 4 w.p.i, 6 w.p.i, 8 w.p.i, and 12 w.p.i. (*Mct1^fl/fl^ n=8*, and *Mct1^iCKO^ n=6*). Significance determined by unpaired two-tailed t-tests and multiple comparisons corrected two-tailed t-tests (j,k).

**Extended Data Figure 7.**
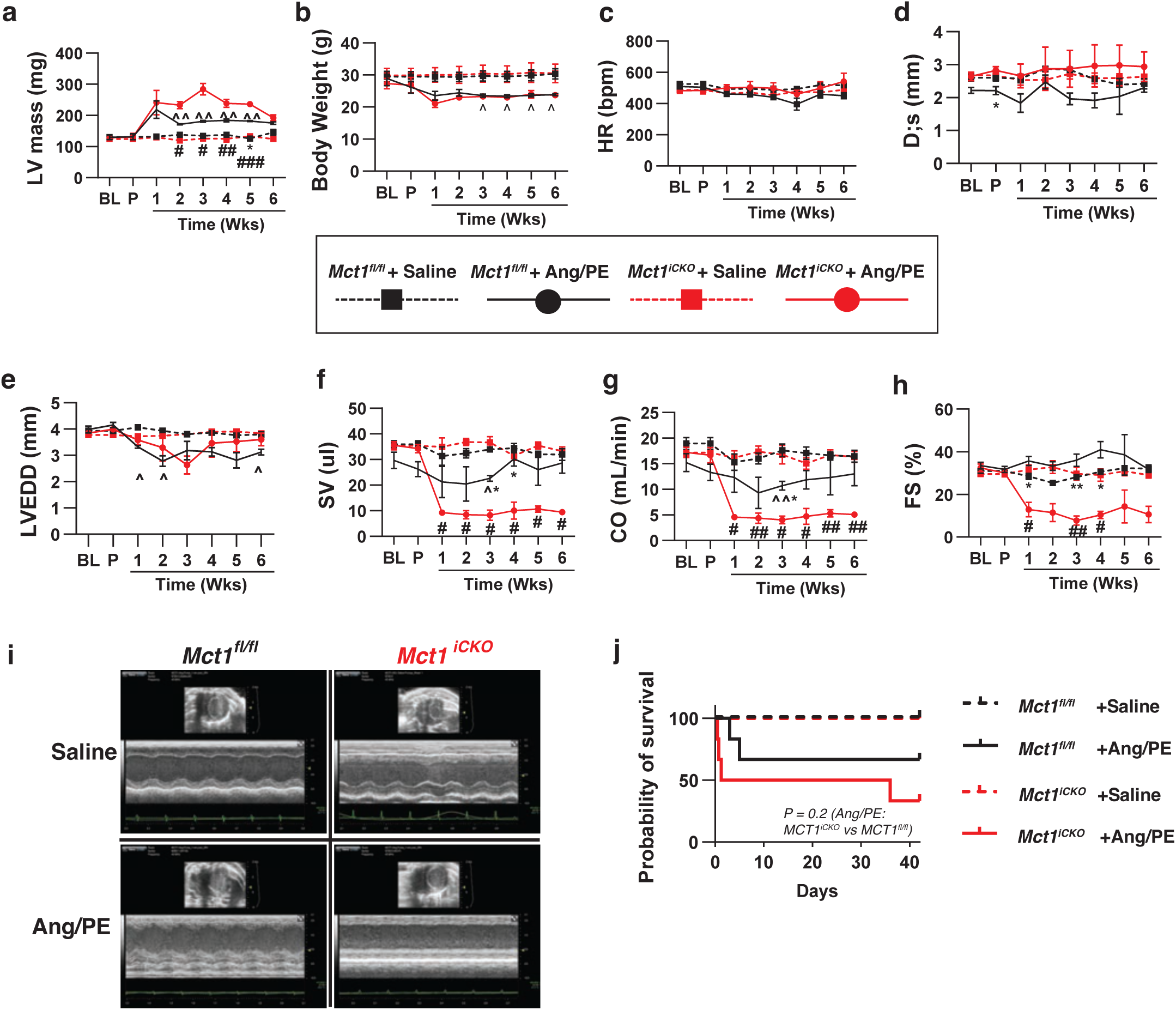
MCT1 deficient hearts have impaired cardiac function upon ANG/PE treatment. Mice (*Mct1^fl/fl^*, and *Mct1^iCKO^*) were treated with angiotensin II (Ang; 1.5 μg/g/day) and phenylephrine (PE; 50 μg/g/day), or normal saline (0.9%, sterile) by osmotic minipump for six weeks (42 days) (*Mct1^fl/fl^* saline *n=7*, *Mct1^fl/fl^* Ang/PE *n=8*, *Mct1^iCKO^*saline *n=4*, *Mct1^iCKO^* Ang/PE *n*=*10*). Cardiac function was assessed by weekly echocardiography. **a**, Absolute left ventricular mass (LV mass: mg). **b**, Body weight (g). **c**, Heart rate (HR: bpm). **d**, Left ventricular end systolic diameter (D;s: mm). **e**, Left ventricular end diastolic diameter (LVEDD: mm). **f**, Stroke volume (SV: μL). **g**, Cardiac output (CO: mL/min). **h**, Fractional shortening (FS: %). **i**, Representative echocardiography images for *Mct1^fl/fl^*, and *Mct1^iCKO^* treated with either saline or Ang/PE. **j**, Kaplan-Meier curve showing probability of survival throughout the six-week administration of Ang/PE in the *Mct1^fl/fl^*, and *Mct1^iCKO^* treated with either saline or Ang/PE. For all echocardiography measurements, a mixed-effects model (repeated measures ANOVA) with the Geisser-Greenhouse correction was used. If significant (p<0.05), then a Tukey’s multiple comparison test was applied with individual variances computed for each comparison. ^=p<0.05, ^^=p<0.01 comparing *Mct1^fl/fl^*Ang/PE vs. *Mct1^fl/fl^* saline. *=p<0.05, **=p<0.01 comparing *Mct1^fl/fl^* Ang/PE vs. *Mct1^iCKO^*Ang/PE. #=p<0.05, ##=p<0.01, ###=p<0.001 comparing *Mct1^iCKO^*Ang/PE vs. *Mct1^iCKO^* saline.

**Extended Data Figure 8.**
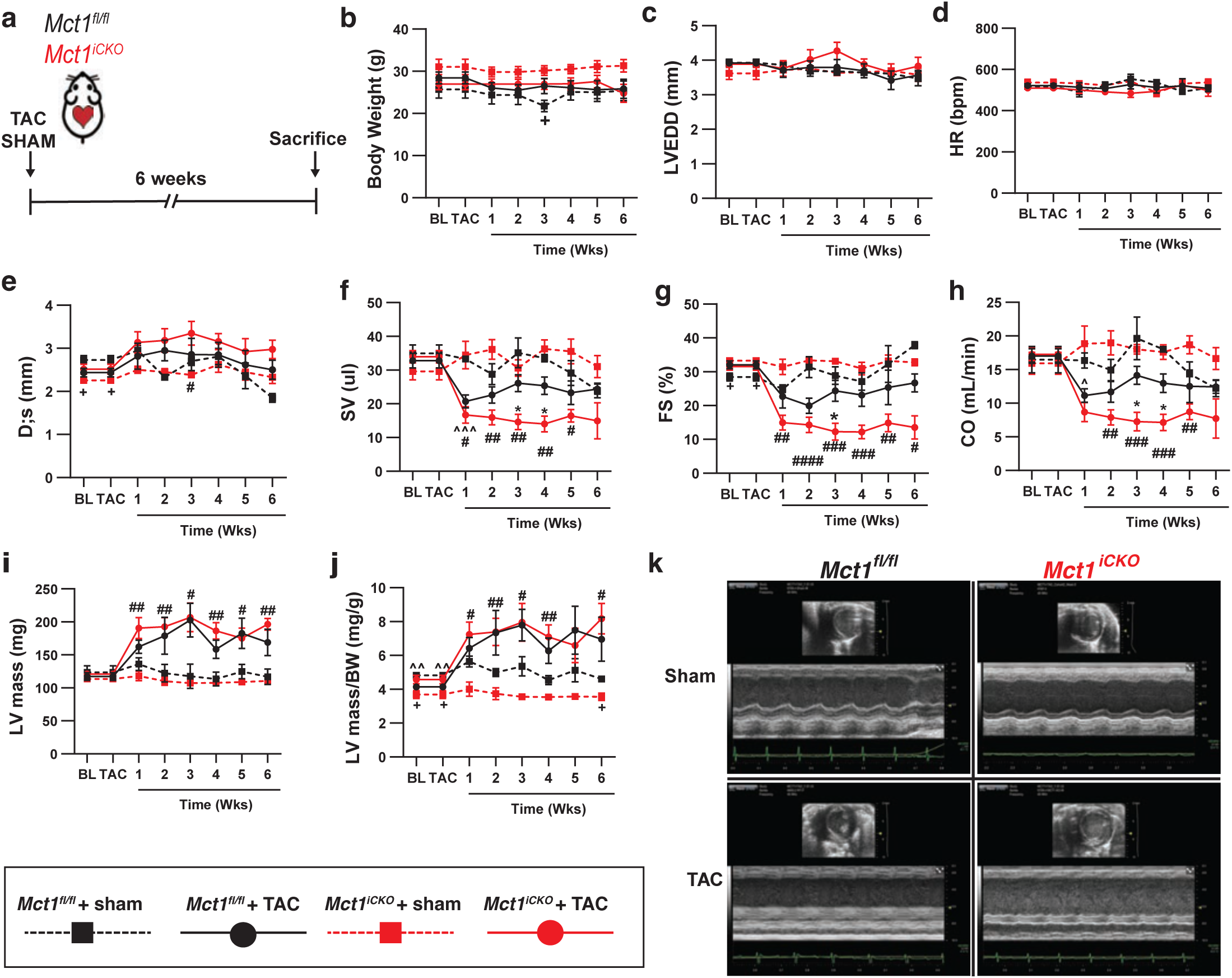
MCT1 deficient hearts have impaired cardiac function upon TAC-induced pressure overload. Mice (*Mct1^fl/fl^*, and *Mct1^iCKO^*) were subjected to transverse aortic constriction, or a sham surgery (*Mct1^fl/fl^* sham *n=5*, *Mct1^fl/fl^* TAC *n=9*, *Mct1^iCKO^* sham *n=4*, *Mct1^iCKO^*TAC *n=10*). Cardiac function was assessed by weekly echocardiography. **a**, Schematic of experimental design. **b**, Body weight (g). **c**, Left ventricular end diastolic diameter (LVEDD: mm). **d**, Heart rate (HR: bpm). **e**, Left ventricular end systolic diameter (D;s: mm). **f**, Stroke volume (SV: μL). **g**, Fractional shortening (FS: %). **h**, Cardiac output (CO: mL/min). **i**, absolute left ventricular mass (LV mass: mg). **j**, Standardized LV mass to body weight (LV mass/BW: mg/g). **k**, Representative echocardiography images for *Mct1^fl/fl^*, and *Mct1^iCKO^* that underwent TAC or sham surgery. A mixed-effects model (repeated measures ANOVA) with the Geisser-Greenhouse correction was used. If significant (p<0.05), then a Tukey’s multiple comparison test was applied with individual variances computed for each comparison. ^=p<0.05, ^^=p<0.01 comparing *Mct1^fl/fl^* TAC vs. *Mct1^fl/fl^* Sham. *=p<0.05, **=p<0.01 comparing *Mct1^fl/fl^* TAC vs. *Mct1^iCKO^* TAC. #=p<0.05, ##=p<0.01, ###=p<0.001 comparing *Mct1^iCKO^*TAC vs. *Mct1^iCKO^* Sham. +=p<0.05, comparing *Mct1^fl/fl^*Sham vs. *Mct1^iCKO^* Sham.

**Extended Data Figure 9.**
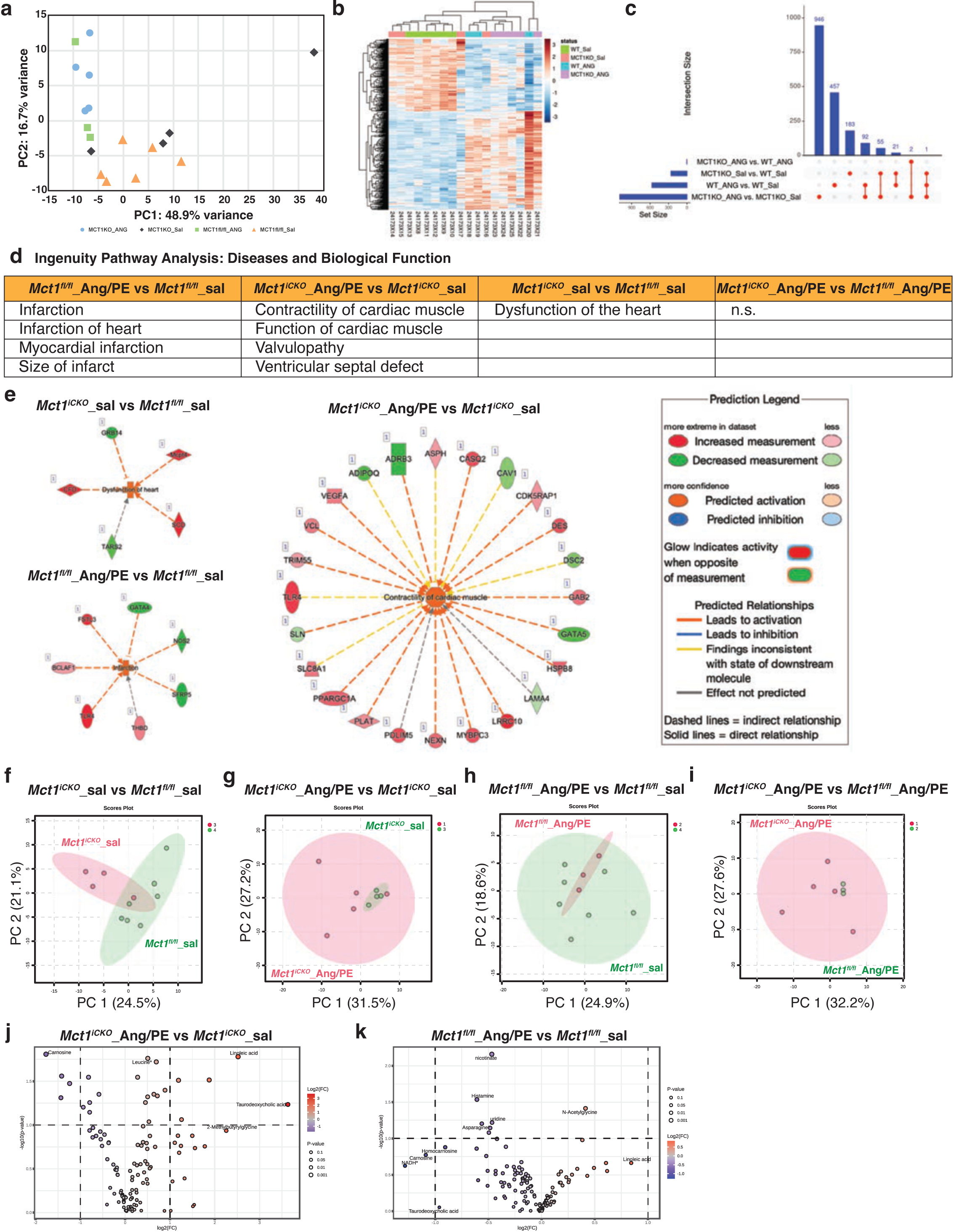
Metabolic and transcriptomic changes upon loss of MCT1. **a,** PCA plot of first two principal components based on the top 500 most variable genes obtained from whole-heart RNA from *Mct1^fl/fl^* and *Mct1^iCKO^*animals treated with either saline or Ang/PE (*Mct1^fl/fl^* saline *n=6*, *Mct1^fl/fl^* Ang/PE *n=3*, *Mct1^iCKO^* saline *n=4*, *Mct1^iCKO^* Ang/PE *n=5*). **b,** Heatmap depicted clustering of samples by 500 most differentially expressed genes. **c,** UpSet plot of all differentially expressed genes between groups. **d,** Ingenuity Pathway Analysis highlighting pathways dysregulated between genotypes within treatment and control groups. **e,** Individual differentially expressed genes and their predicted contributions to pathways “infarction”, “dysfunction of heart”, and ‘contractility of cardiac muscle” from *Mct1^fl/fl^* and *Mct1^iCKO^* animals treated with either saline or Ang/PE. **f-i,** PCA plot of the first two principal components based on all quantified polar metabolites from the hearts from *Mct1^fl/fl^* and *Mct1^iCKO^* animals treated with either saline or Ang/PE. Plotted are all pairwise comparisons using the same tissue samples as RNAseq data in a (*Mct1^fl/fl^* saline *n=6*, *Mct1^fl/fl^* Ang/PE *n=3*, *Mct1^iCKO^* saline *n*=4, *Mct1^iCKO^* Ang/PE *n*=5). **j,k,** Volcano plots showing statistically significantly differentially abundant metabolites from hearts of Ang/PE and saline treated *Mct1^iCKO^* and *Mct1^fl/fl^* animals respectively (*Mct1^fl/fl^* saline *n=7*, *Mct1^fl/fl^* Ang/PE *n=3*, *Mct1^iCKO^* saline *n=4*, *Mct1^iCKO^* Ang/PE *n=5*).

**Extended Data Figure 10.**
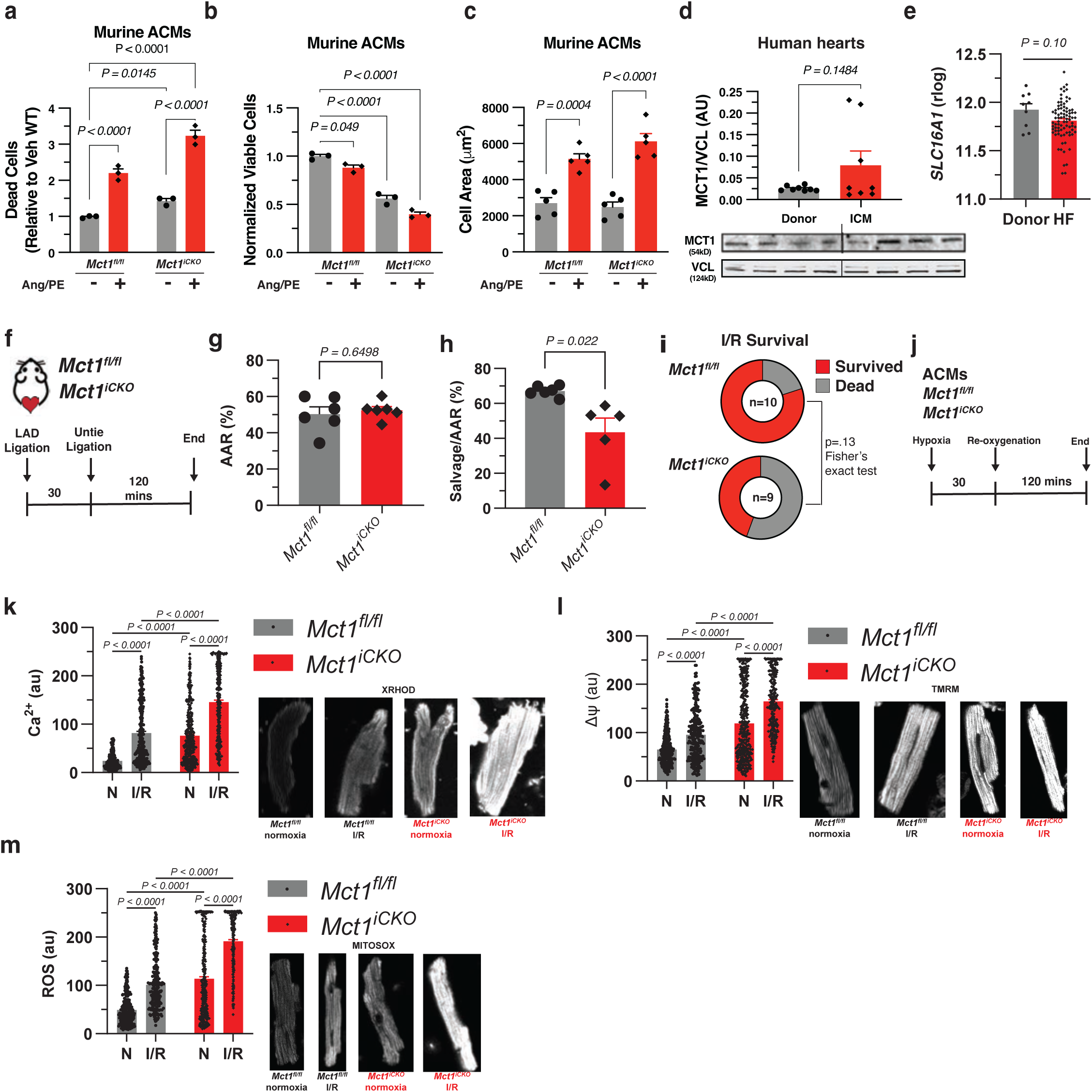
*Mct1* deletion leads to enhanced cell death, increased hypertrophy, and impairs mitochondrial function following myocardial stress. **a**, Number of dead cells (relative to vehicle treated *Mct1^fl/fl^* cardiomyocytes) following administration of angiotensin II and phenylephrine (Ang/PE) or vehicle (Veh) to primary adult cardiomyocytes (ACMs) from *Mct1^fl/fl^*and *Mct1^iCKO^* hearts (*n=3*). **b**, Normalized cell viability after Ang/PE or Veh treatment in ACMs from *Mct1^fl/fl^*and *Mct1^iCKO^* (*n=3*). **c**, Cell area (μm^2^) of *Mct1^fl/fl^* and *Mct1^iCKO^*ACMs treated with Ang/PE or Veh (*n=5*). **d**, Relative MCT1 abundance normalized to VDAC (AU: arbitrary units) in non-failing donor heart samples compared to samples acquired from advanced heart failure (HF) patients undergoing transplant due to ischemic cardiomyopathy (ICM) (donors *n=8*, ICM *n=6*). **e**, Standardized r-log values from transcriptomics (RNA sequencing) comparing non-failing donor heart samples to samples acquired from advanced HF patients at the time of left ventricular assist device (LVAD) therapy. **f**, Schematic of in vivo acute ischemia-reperfusion injury (I/R) in *Mct1^fl/fl^*, and *Mct1^iCKO^* mice. **g**, Area at risk (AAR: %) of I/R injury (*n=6*). **h**, Myocardial salvage standardized to the AAR (*n=6*). **i**, Survival rates of the mice subjected to acute I/R injury (*Mct1^fl/fl^ n=10*, *Mct1^iCKO^ n=13*). **j**, Schematic of in vitro hypoxia and reoxygenation experiment using ACMs from *Mct1^fl/fl^*, and *Mct1^iCKO^* hearts. **k-m**, Mitochondrial specific reporters for calcium (Ca^2+^: XRhod), membrane potential (Δψ: TMRM), and reactive oxygen species (ROS: mitosox), respectively in ACMs isolated from *Mct1^fl/fl^*, and *Mct1^iCKO^* hearts. Significance determined by two-tailed unpaired t tests (d-h) and one-way and two-way ANOVA (a-c, k-m) with Tukeys HSD multiple comparison test. ns, not significant (p>0.05).

**Extended Data Figure 11.**
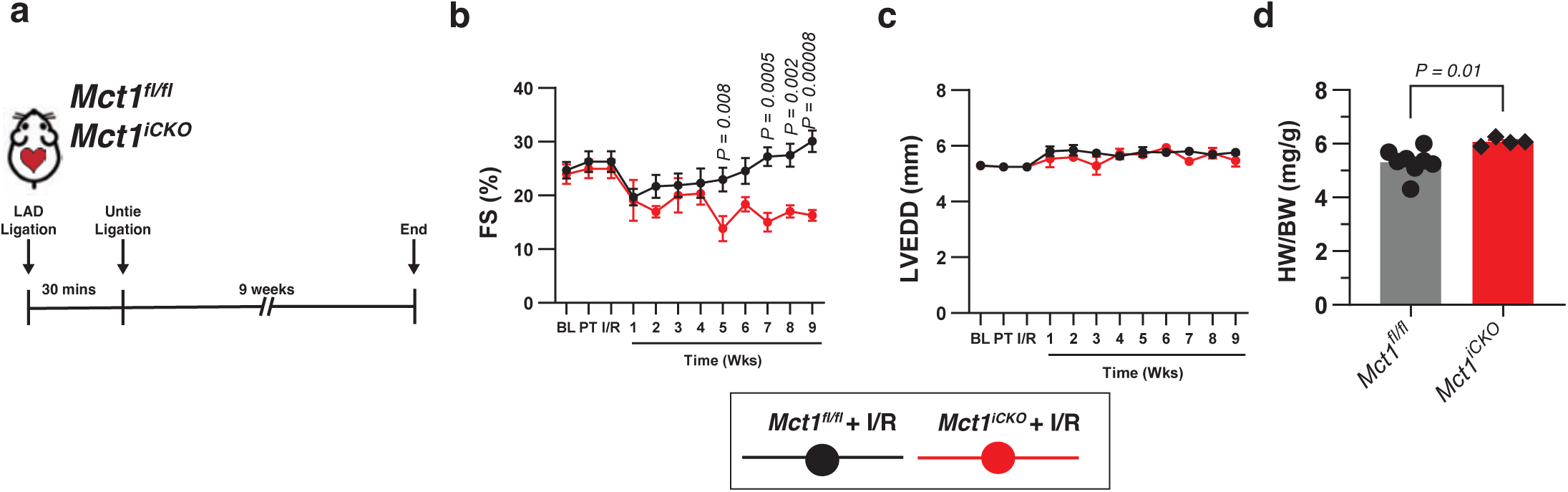
Loss of MCT1 impairs recovery from myocardial ischemia reperfusion injury. **a**, Diagram representing in vivo chronic ischemia-reperfusion injury (I/R) in *Mct1^fl/fl^*, and *Mct1^iCKO^* mice. Mice were subjected to cardiac injury and surviving mice allowed to recover (*Mct1^fl/fl^ n=11*, and *Mct1^iCKO^ n=9*). Echocardiograms were performed weekly for nine weeks following I/R injury. **b**, Fractional shortening (FS: %). **c**, Left ventricular end diastolic diameter (LVEDD: mm). **d**, Heart weight standardized to body weight (mg/h) in *Mct1^fl/fl^*, and *Mct1^iCKO^* mice at the end of nine-weeks following I/R injury. Data are plotted as mean ± SEM. Significance determined by unpaired t-two tailed t-test with multiple comparisons correction. ns, not significant (p>0.05).

