## Supplemental Data A for "Direct mitochondrial import of lactate supports resilient carbohydrate oxidation"

### Slide 1
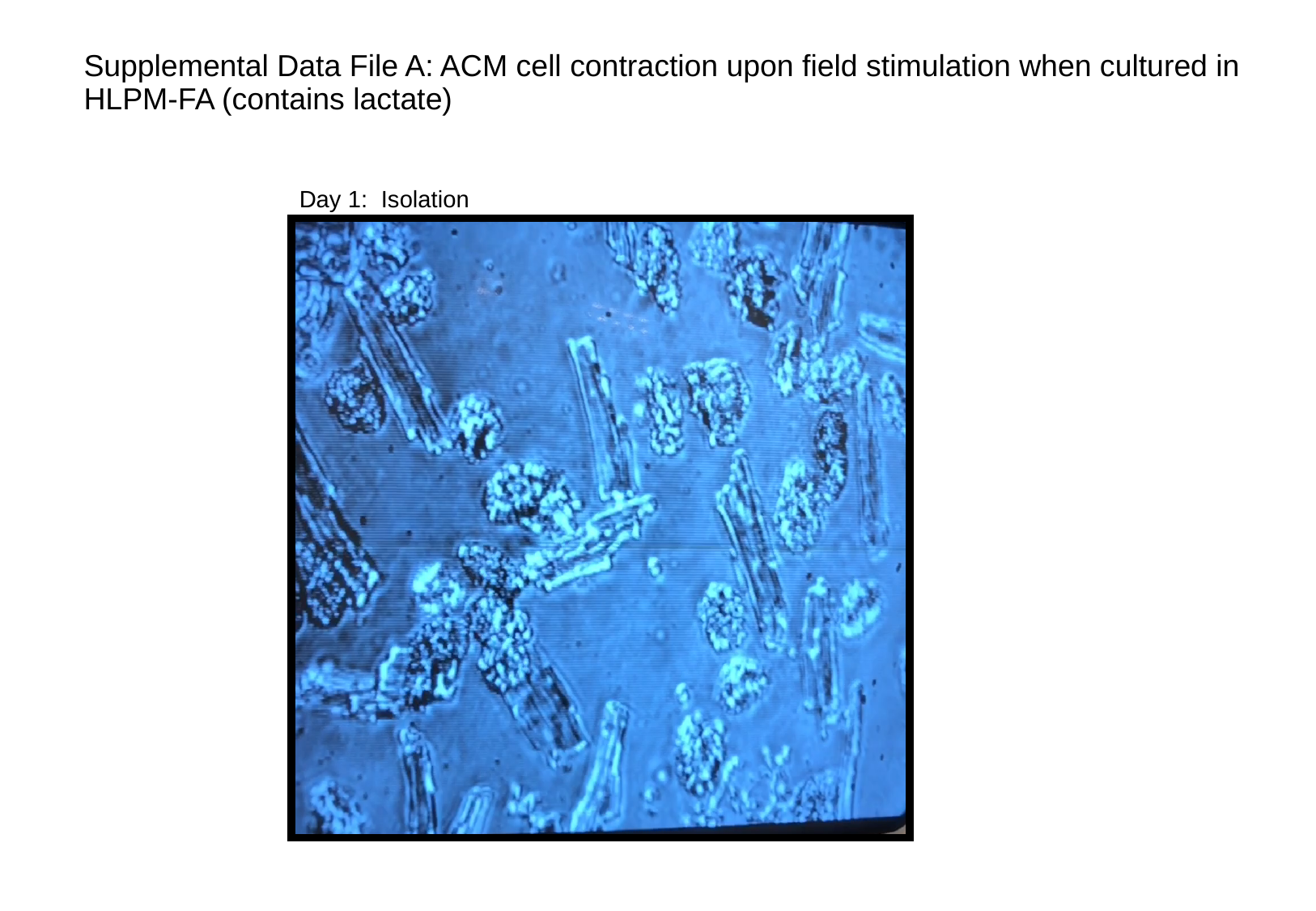

#### Supplemental Data File A: ACM cell contraction upon field stimulation when cultured in HLPM-FA (contains lactate)
Day 1: Isolation

### Slide 2
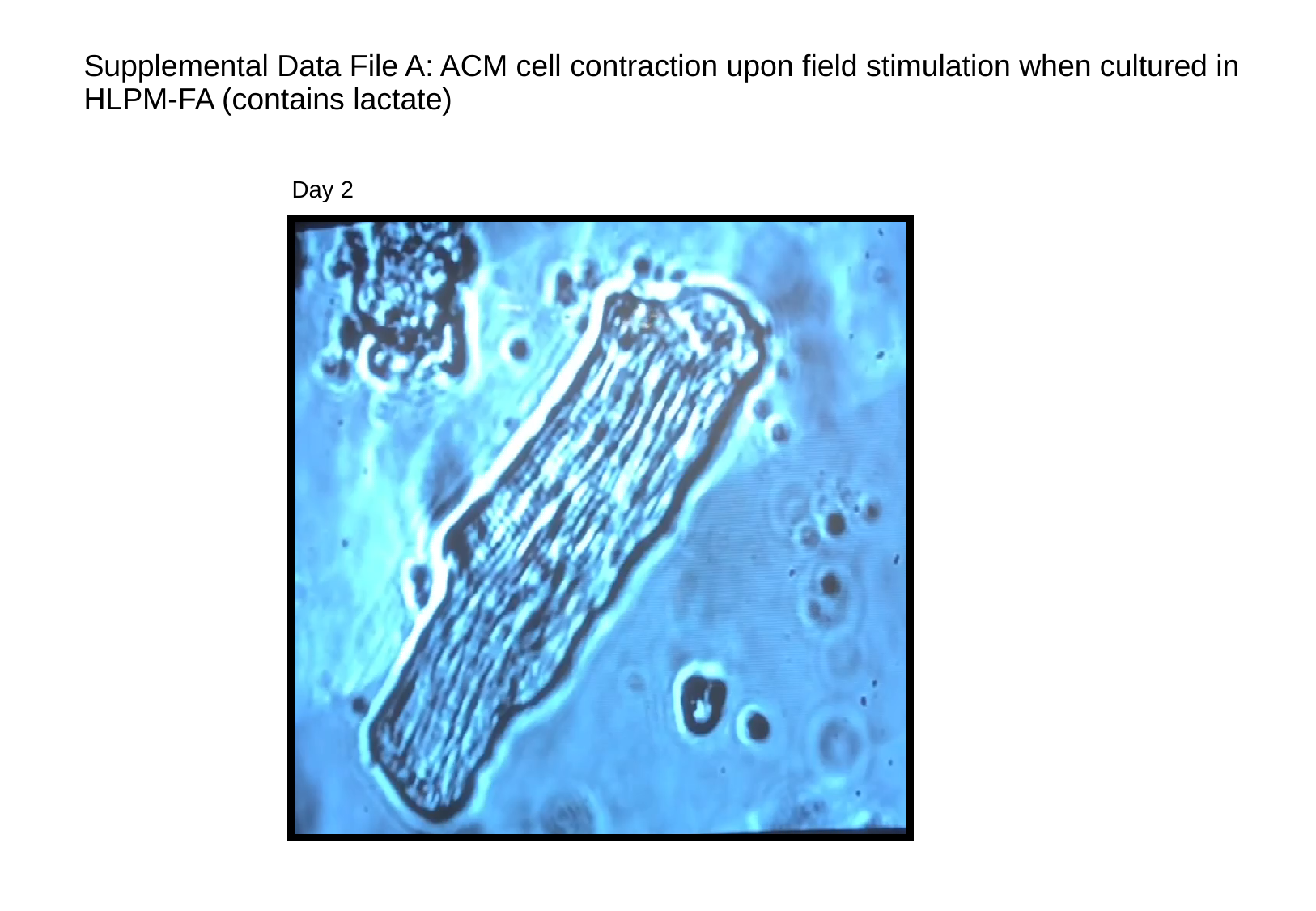

#### Supplemental Data File A: ACM cell contraction upon field stimulation when cultured in HLPM-FA (contains lactate)
Day 2

### Slide 3
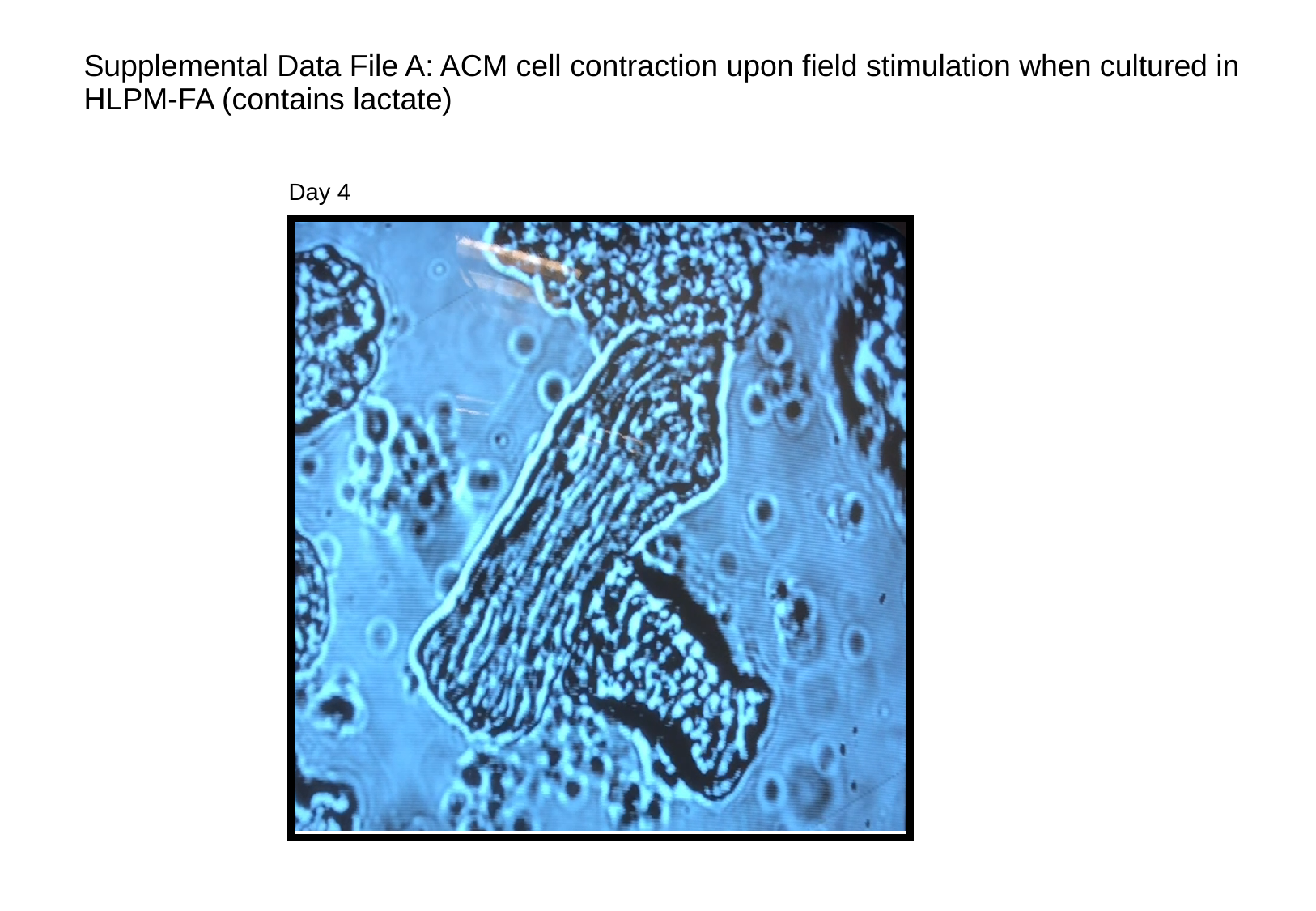

#### Supplemental Data File A: ACM cell contraction upon field stimulation when cultured in HLPM-FA (contains lactate)
Day 4

### Slide 4
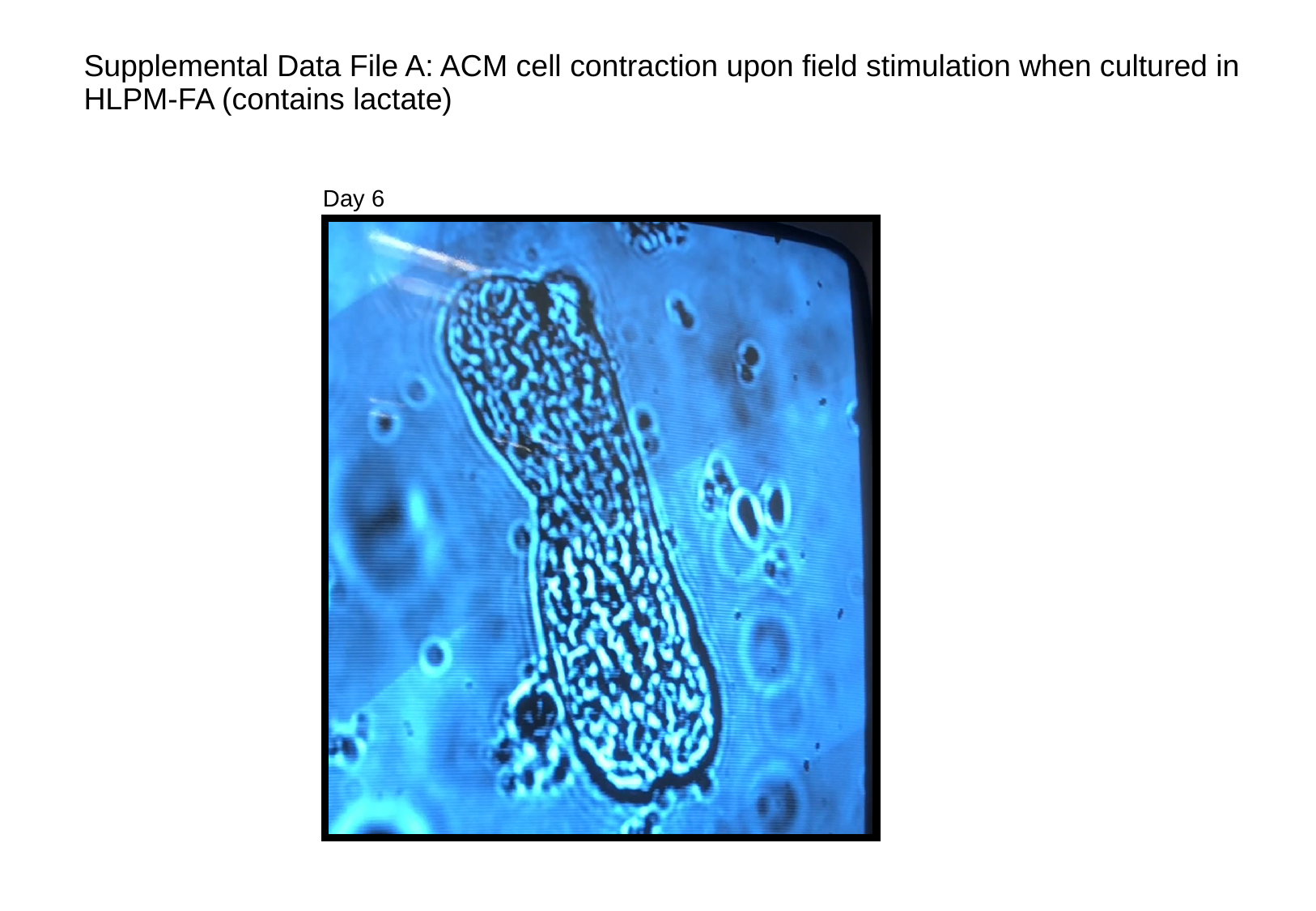

#### Supplemental Data File A: ACM cell contraction upon field stimulation when cultured in HLPM-FA (contains lactate)
Day 6
